# Phosphoproteome-derived peptide libraries for deep specificity profiling of phosphatases and phospholyases

**DOI:** 10.1101/2025.08.12.669978

**Authors:** Katarzyna Radziwon, Laura A. Campbell, Lauren E. Mazurkiewicz, Sopo Jalalishvili, Izabelle Eppinger, Aanika Parikh, Amy M. Weeks

## Abstract

Protein phosphorylation is dynamically regulated by the opposing activities of phosphowriter enzymes (kinases) and phosphoeraser enzymes (phosphatases and phospholyases). While significant progress has been made toward defining the sequences preferences of kinases, the selectivity of phosphoerasers has not been explored at scale. Here, we develop an experimental platform based on tandem mass spectrometry analysis of phosphoproteome-derived peptide libraries (PhosPropels) to map phosphoeraser activity across thousands of biologically relevant phosphosites. We extract positional residue preferences to rapidly define sequence motifs recognized by eight phosphoerasers spanning diverse species of origin, protein folds, and enzymatic mechanisms. Taking advantage of the throughput of our approach, we profiled 34 variants of the phosphothreonine lyase OspF from *Shigella flexneri*, uncovering an intrinsic preference for p38 and Erk MAP kinase activation loops and revealing the enzyme residues that influence its selectivity for phosphothreonine. Our results establish a general method for linking phosphorylation sites to the enzymes that remove them, providing a means to dissect a key component of cellular regulatory networks.

## Main text

Protein phosphorylation is a dynamic post-translational modification that regulates nearly every aspect of biology^1,2^. Phosphorylation signaling can be understood in terms of a well-established read/writer/eraser framework: kinases write phosphorylation marks, reader proteins (or intramolecular sensing mechanisms) interpret them to direct downstream signaling, and erasers (such as phosphatases and phospholyases) remove them to regulate signaling dynamics^3,4^. While significant progress has been made in understanding the specificity of kinases^5,6^ and reader domains^7,8^, much less is known about phosphoeraser specificity. This gap stems largely from the challenges and costs of phosphopeptide synthesis and the need to obtain precise positional information.

Synthetic phosphopeptide-based approaches present challenges for the high-throughput analysis that is needed for comprehensive phosphoeraser substrate selectivity profiling^9^. Array-based or barcoded library approaches rely on individually addressable peptides that are synthesized one by one and typically use binary readouts of phosphorylation, precluding analysis of multiply phosphorylated peptides^10–12^. Pooled libraries offer greater synthetic scalability but are limited in size by the constraints of typical MS analysis workflows, requiring researchers to choose which motifs and variations to include in the library^13^. Higher throughput approaches that query phosphoeraser sequence selectivity in the context of diverse, biologically relevant substrate pools are therefore needed to advance our understanding of phosphorylation signaling networks.

An optimal method for deep profiling of phosphoeraser substrate sequence specificity would (1) enable multiplexed analysis of a proteome-scale pool of biologically relevant substrates; (2) reveal the exact site(s) of dephosphorylation in singly or multiply phosphorylated peptides; and (3) rely on an easily accessible source of diverse phosphopeptides. Inspired by proteomic approaches for probing protease and ligase specificity^14–16^, we developed a tandem mass spectrometry (LC-MS/MS)-based *in vitro* assay for dephosphorylation of peptides derived from the human phosphoproteome. Our method uses statistical comparison of phosphopeptide sequence features in enzyme-treated samples versus untreated controls to generate quantitative specificity profiles for phosphoeraser enzymes. These profiles reflect position-specific substrate preferences across thousands of biologically derived phosphopeptides, including features like adjacent phosphosites that are often overlooked when using synthetic peptide libraries.

We applied our method to extract positional residue preferences for five phosphatases and three phosphothreonine lyases derived from phage, human, and bacteria, highlighting its utility for analyzing enzymes with diverse species of origin, protein folds, enzymatic mechanisms, and molecular functions. Leveraging the throughput of our approach, we profiled 20 variants of the phosphothreonine (pThr) lyase OspF from *Shigella flexneri* to define the molecular basis of its substrate specificity. We find that OspF’s catalytic domain has an intrinsic preference for the pThr-Xaa-pTyr (where Xaa ≠ Pro) motif found in p38 and Erk MAP kinase (MAPK) activation loops, independent of its MAPK docking motif. We also observe that residues that do not directly contact the substrate have a strong influence on OspF’s selectivity for pThr over phosphoserine (pSer) sites. Together, our results provide significant insights into phosphoeraser specificity and establish a general method for querying phosphoeraser specificity at proteome scale.

## Results

### Phosphoproteome-derived peptide libraries (PhosPropels) encompass biologically relevant phosphoeraser substrates

To obtain a diverse, biologically relevant pool of substrates for phosphoeraser profiling, we generated phosphoproteome-derived peptide libraries (PhosPropels) from HEK293T cells (**Fig. 1a, b**)^17^. Tryptic digestion of protein extracts followed by TiO_2_ enrichment yielded 9,012±767 unique phosphopeptides per replicate, which comprised 90±5% of total peptides identified (**Fig. 1c**, **Supplementary Dataset 1**). These peptides harbored 9,888±989 high-confidence phosphorylation sites (ptmRS site localization probability >90%, **Fig. S1**) per replicate whose composition was consistent with previous measurements of the human phosphoproteome (90.3±0.3% phosphoserine (pSer), 9.1±0.1% phosphothreonine (pThr), and 0.6±0.2% phosphotyrosine (pTyr) residues) (**Fig. 1c**, **Fig. S1**)^18^. To increase representation of pTyr-containing peptides, we pre-treated cells with pervanadate, a broad spectrum pTyr phosphatase inhibitor^19^. This resulted in an approximately 20-fold increase in the number of high-confidence pTyr sites (1,276±346) compared to the untreated library (58±26), expanding its utility for profiling phosphoerasers that recognize pTyr residues (**Fig. 1c**, **Fig. S2**, **Supplementary Dataset 2**). Similar results were obtained when PhosPropels were generated from Jurkat or K562 cells (**Extended Data Fig. 1**, **Fig. S3**, **Fig. S4**, **Supplementary Datasets 3-4**), demonstrating the generalizability of the approach across cell lines.

**Figure 1.**
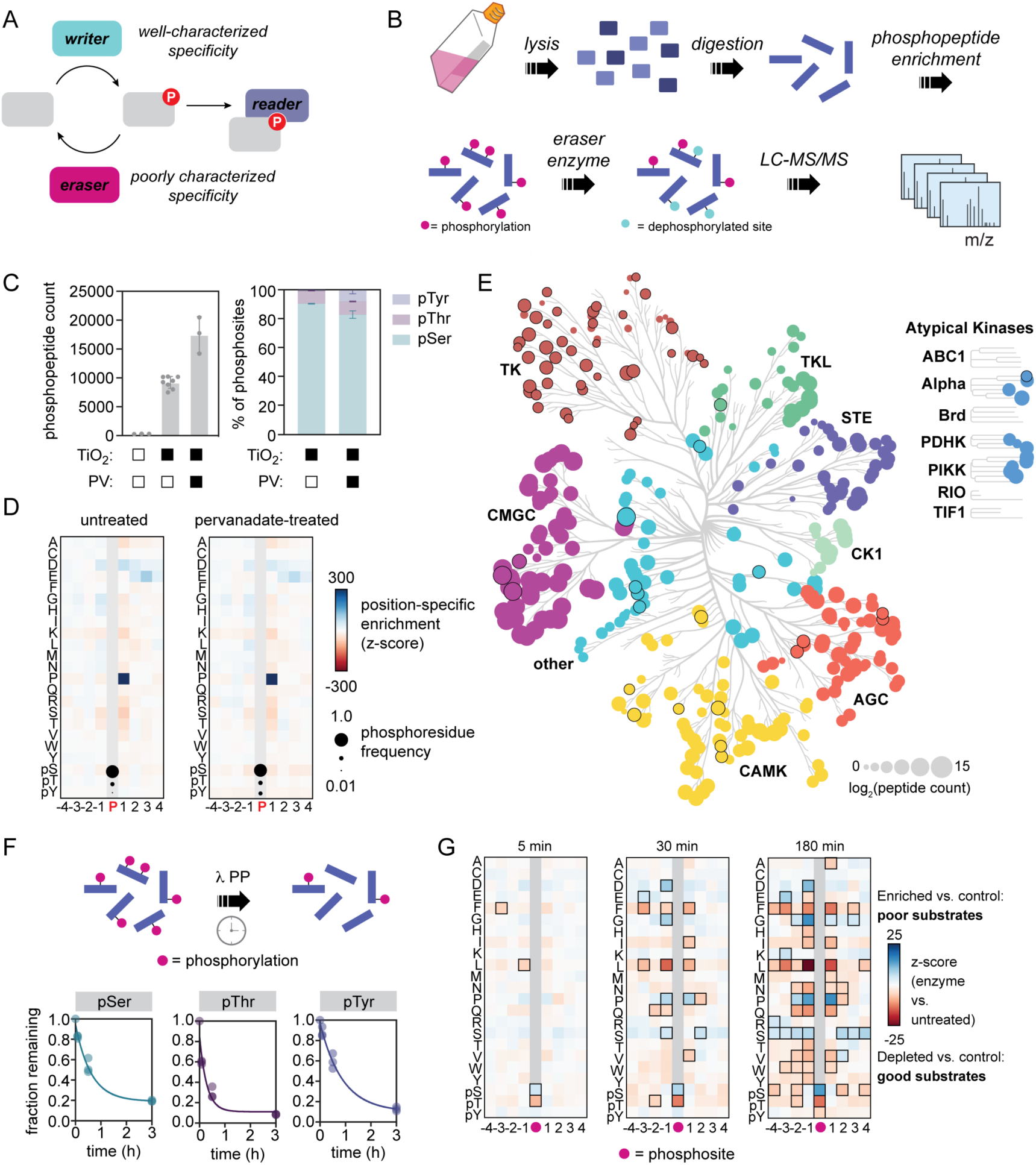
Phosphoproteome-derived peptide libraries (PhosPropels) for probing phosphoeraser specificity at scale. a) Signaling via phosphorylation can be understood in terms of a reader/writer/eraser framework. b) Phosphopeptides enriched from biological samples provide a proteome-scale substrate library for profiling phosphoeraser specificity using LC-MS/MS. c) Titanium dioxide (TiO_2_) enrichment of enables identification of thousands of phosphopeptides from HEK293T cells in a single LC-MS/MS experiment; the distribution of pSer, pThr, and pTyr sites in the sample can be altered by treatment with phosphatase inhibitors such as the pTyr phosphatase inhibitor pervanadate (PV). d) LC-MS/MS analysis of phosphopeptides enables precise localization of phosphorylation modifications and enables analysis of the identity of flanking residues, which include all proteinogenic amino acids, pSer, pThr, and pTyr in each position. e) Phosphopeptides in the library are a biologically relevant substrate pool and contain sequences targeted by all major families of human kinases. Node size represents the average log_2_ count of sequences that score at least 2 standard deviations above the kinase-specific PSSM mean. Nodes with a significant (p < 0.05) increase in sequence count upon pervanadate treatment are outlined in black. f) Treatment of phosphoproteome-derived peptide libraries leads to depletion of pSer, pThr, and pTyr sites. g) Heatmap showing positional enrichment or depletion of amino acid residues flanking phosphosites. Position-residue combinations outlined in black were significantly (p < 0.0001) enriched or depleted compared to an untreated control.

To characterize the diversity of flanking sequences in our libraries, we aligned all high-confidence phosphosites and calculated the position-specific frequency of each amino acid in positions −4 to +4 relative to the phosphosite. We then compared these frequencies to the overall amino acid frequencies in the library by calculating z-scores, enabling detection of position-specific enrichment and depletion (**Fig. 1d**, **Supplementary Note 1**). All 20 proteinogenic amino acids, as well as pSer, pThr, and pTyr, were represented at every position in this window (**Fig. S5**). Most position-specific frequencies reflected overall amino acid abundance in the library; however, Pro at +1 was substantially enriched compared to library abundance. This likely reflects the activity of Pro-directed kinases including cyclin-dependent kinases (CDKs) and mitogen-activated protein kinases (MAPKs), which are active in proliferating cells^20,21^. These data establish a statistical background for downstream profiling of phosphoeraser specificity.

To evaluate the biological diversity of substrates in our libraries, we scored each phosphosite using recently published position-specific scoring matrices for 396 kinases comprising 87% of the active human kinome (**Supplementary Dataset 5**)^5,6^. Untreated HEK293T libraries had robust representation of predicted kinase substrates from all Ser/Thr kinase branches and from the Alpha, PIKK, and PDHK families of atypical kinases, while pervanadate treatment increased representation of tyrosine kinase substrates (**Fig. 1e**). Similar results were obtained with libraries from Jurkat and K562 cells (**Extended Data Fig. 1**). These results show that although PhosPropels sample a relatively small fraction of combinatorial sequence space, they capture a broad array of biologically relevant substrates.

### Statistical profiling of phosphoeraser specificity using PhosPropels

To assess the utility of PhosPropels for profiling phosphoeraser specificity, we treated them with *λ* phosphatase (*λ*PP), a broad specificity enzyme that is often used for global dephosphorylation^22,23^. We quenched reactions at multiple timepoints and analyzed samples by LC-MS/MS to quantify the frequency of pSer, pThr, and pTyr among the remaining phosphorylation sites (**Fig. 1f**) and the position-specific frequency of each amino acid flanking those sites (**Fig. 1g**). We hypothesized that substrates efficiently dephosphorylated by *λ*PP would be rapidly depleted from the library, while poor substrates would persist, allowing sequence prefences to be inferred statistically from depletion and enrichment patterns across thousands of phosphopeptides (**Supplementary Note 1**).

*λ*PP had robust activity on PhosPropels, reducing phosphopeptides from 90% to 14±1 % of total peptides over 3 h (**Fig. 1f**, **Supplementary Dataset 6**). In agreement with previous studies^24^, pThr sites (t_1/2_ = 12±3 min) were depleted from the library faster than pSer sites (t_1/2_ = 26±4 min) and pTyr sites (t_1/2_ = 36±6 min), and phosphosites with Pro at the +1 position were strongly disfavored as substrates (**Fig. 1f,g**). We also identified previously unrecognized *λ*PP substrate preferences: Pro was disfavored at the −1 and −2 positions, while Phe, Leu, and Gln were preferred at the +1 and −1 positions. pSer, pThr, and pTyr were depleted at all postions outside the central phosphosite, consistent with the ability of *λ*PP to act on multiple phosphosites within the same peptide. These results demonstrate that our method accurately reports on phosphatase substrate specificity, recapitulating known trends and revealing new ones.

### PhosPropels for profiling the selectivity of human pTyr and pSer/pThr phosphatases

We next profiled the catalytic domain of the human pTyr phosphatase PTP1B (residues 1-321), an important regulator of metabolic and oncogenic signaling pathways that is mechanistically distinct from *λ*PP (**Fig. 2a**)^25,26^. We treated PhosPropels (0.1 mg/mL) with PTP1B_1-321_ (20 nM), quenched the reactions at various timepoints, and analyzed the position-specific enrichment and depletion of each amino acid surrounding the remaining phosphosites. PTP1B_1-321_ rapidly depleted pTyr, but not pSer or pThr, from the library, consistent with both its well-established pTyr specificity and measurements of phosphate release from individual synthetic peptides (**Fig. 2b**, **c**, **d**, **Fig****. S6**, **Supplementary Dataset 7**). We therefore focused our analysis on features flanking pTyr sites.

**Figure 2.**
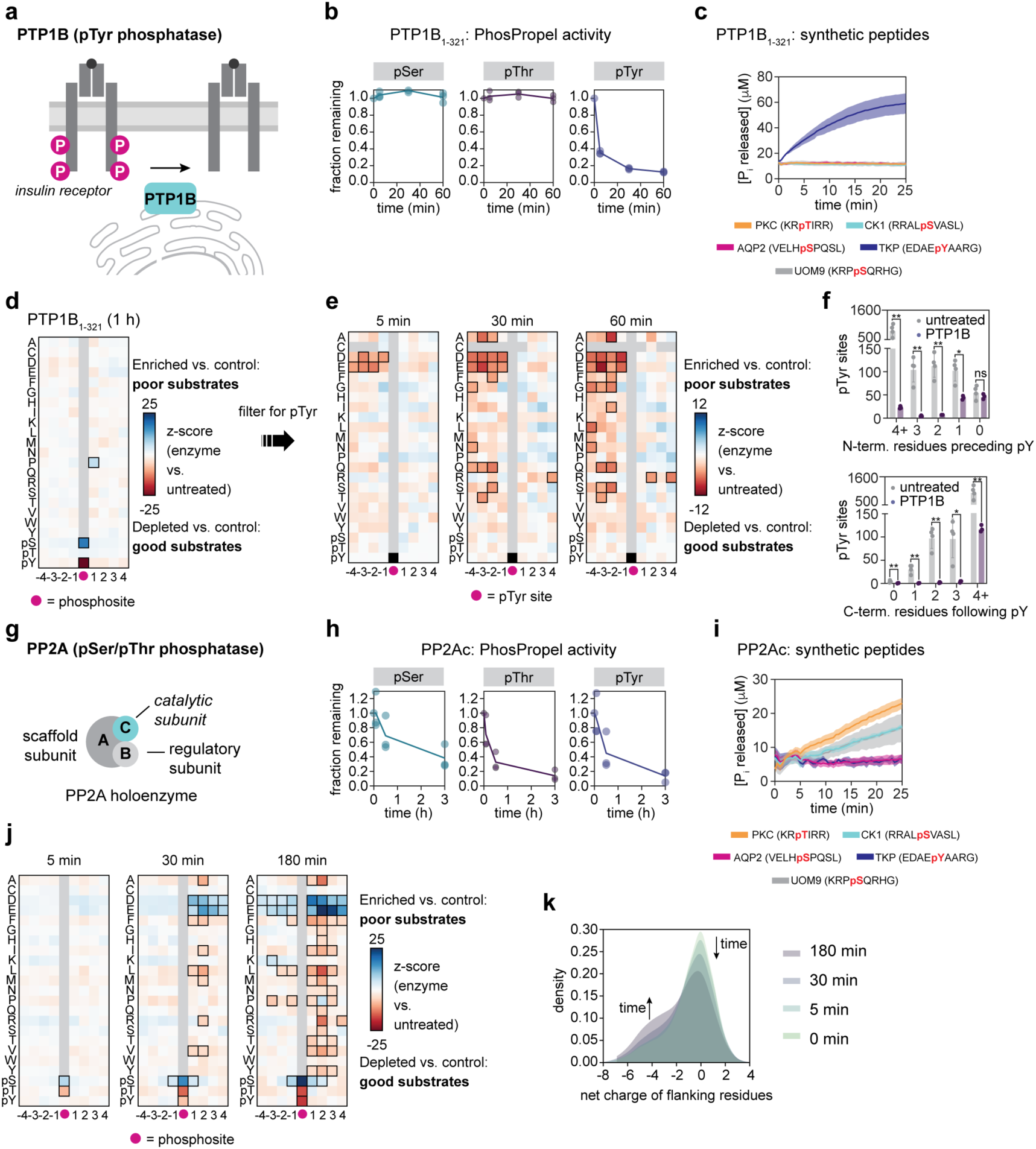
Deep profiling of human phosphatase specificity using PhosPropels. a) Human PTP1B is protein tyrosine phosphatase that regulates metabolic signaling. b) PTP1B_1-321_ rapidly depletes pTyr sites, but not pSer or pThr sites, from PhosPropels Data show phosphosite counts from n = 3 biological replicates. c) PTP1B_1-321_-catalyzed phosphate release from a panel of synthetic human phosphopeptides Line shows average of n=3 biological replicates and filled area shows s.d. d) PTP1B_1-321_ significantly depletes pTyr from the PhosPropel, while pSer sites and sites with Pro at +1 become significantly enriched. e) Among pTyr sites, PTP1B_1-321_ favors those with acidic residues on the N-terminal side. f) Top, pTyr sites with fewer than two amino acids on the N-terminal side persist in the library after 2 h of PTP1B_1-321_ treatment. Bottom, PTP1B_1-321_ does not require additional amino acids following pTyr on the C-terminal side. Data represent pTyr counts from n = 3-5 biological replicates. pTyr counts in the treated vs. untreated samples were compared using unpaired *t*-tests and p-values were corrected for multiple comparisons using the Holm-Šídák method. * p < 0.05; ** p < 0.01; ns, not significant. g) PP2Ac is the catalytic subunit of the pSer/pThr phosphatase PP2A. h) PP2Ac depletes pSer, pThr, and pTyr sites from PhosPropels. Data show phosphosite counts from n = 3 biological replicates. i) PP2Ac-catalyzed phosphate release from a panel of synthetic human phosphopeptides. Line shows average of n=3 biological replicates and filled area shows s.d. j) PP2Ac disfavors substrates with flanking acidic residues. k) Kernel density estimate plot showing that phosphosites whose flanking residues have a high net negative charge accumulate in the library over time. For panels (d), (e), and (j), z-scores were calculated by comparing positional frequencies to the 0 min timepoint using counts summed across n = 3 biological replicates. Residue-position combinations with Benjamini-Hochberg FDR-adjusted p-values < 0.0001 are outlined in black.

Consistent with previous studies, our data revealed strong sequence preferences on the N-terminal side of the pTyr site (**Fig. 2e**)^27,28^. Within 5 min, pTyr sites with acidic residues in the −1 to −4 positions were substantially depleted from the library. After 30 min, pTyr sites with Gln at positions −2 to −4 were also depleted. At the 60 min timepoint, the majority of pTyr sites (88±1%) were depleted. Most (72±3%) pTyr sites remaining after 60 min had the pTyr residue within two amino acids of the N terminus, highlighting the importance of N-terminal amino acids in substrate recognition (**Fig. 2f**). This feature was uniquely identifiable using our method, which incorporates phosphopeptides of variable length.

In contrast, C-terminal sequence preferences were weaker. Only pTyr sites with Arg at the +2 or +4 positions were significantly depleted among the pTyr sites remaining after 60 min (**Fig. 2e**), suggesting that PTP1B tolerates a broad array of C-terminal residues. Many of PTP1B’s best studied substrates, including insulin receptor, contain a tandem pTyr motif (pYpY)^29^. While pYpY sites with pTyr at the +1 position were rapidly depleted from the library upon PTP1B treatment, depletion was not statistically significant (p > 0.05), likely due to the relatively small number of pYpY sites in the library. However, we note that the pYpY site in insulin receptor (^1158^pYETD<u>pYpY</u>RKGG^1167^) is flanked by other features we found to be favorable (**Fig. 2e**), including acidic residues on the N-terminal side and Arg at the +2 position.

We next profiled the catalytic subunit of PP2A (PP2Ac), a pSer/pThr phosphatase involved in apoptosis, metabolism, and cell migration^30,31^. In cells, PP2Ac assembles with a scaffold A subunit and regulatory B subunits that direct its substrate selectivity (**Fig. 2g**). A complete understanding of how PP2Ac’s activity is shaped by these protein partners requires detailed knowledge of its intrinsic specificity prior to maturation and holoenzyme formation. To gain this understanding, we profiled PP2Ac using PhosPropels. PP2Ac exhibited broad dephosphorylation activity, depleting pSer, pThr, and pTyr from the library over 3 h (**Fig. 2h**, **Supplementary Dataset 8**). Analysis of phosphate release from individual peptides confirmed PP2Ac’s broad substrate specificity (**Fig. 2i**), including previously observed basal pTyr activity^32^, which was not captured in a recent large-scale PP2Ac profiling study as pTyr sequences were not included in the library design^13^.

Analysis of flanking residues preferred by PP2Ac broadly agreed with previous specificity studies using synthetic pSer/pThr peptides (**Fig. 2j**, **Fig. S7**). Phosphosites with Pro at the +1 or +2 positions or with acidic residues on the C-terminal side were poor substrates, while those with aromatic or hydrophobic residues at +1 and +2 were preferred. There was also a preference for Arg or Ala at the +2 position. Supporting these observations, phosphopeptides with large net negative charges persisted in the library following PP2Ac treatment (**Fig. 2k**). These results demonstrate that PP2Ac can discriminate among diverse biological substrates based on its active site properties, even in the absence of regulatory subunits, and lay the groundwork for future studies exploring how holoenzyme formation shapes substrate selectivity.

### PhosPropels reveal unique substrate profiles of related bacterial effector phosphatases

Bacterial pathogens have evolved effector enzymes that modulate host phosphorylation signaling to promote infection (**Fig. 3a**)^33,34^. We applied the PhosPropel approach to two phosphatase effectors from *Legionella pneumophila,* WipA and WipB, whose specificity profiles remain incompletely characterized^35–37^. Both enzymes contain phosphoprotein phosphatase (PPP) family phosphatase domains, but WipA has been classified as a pTyr phosphatase that inhibits F-actin polymerization and WipB has been classified as a pSer/pThr phosphatase that targets proteins involved in lysosomal nutrient sensing. These assignments were based on limited *in vitro* assays on individual peptides. To better define their specificities, we profiled their activity on PhosPropels (**Fig. 3b**).

**Figure 3.**
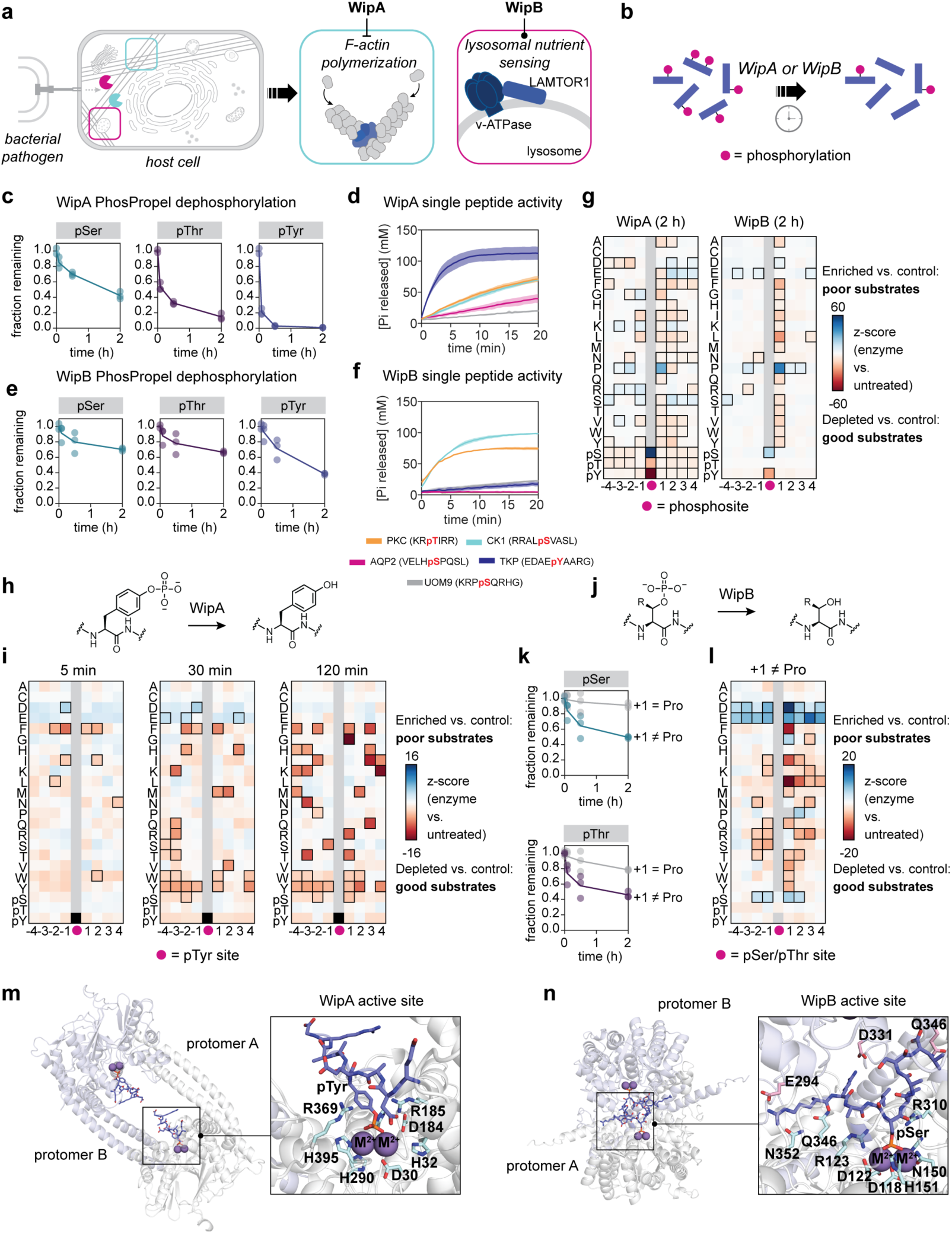
PhosPropels reveal unique substrate profiles of bacterial effector phosphatases WipA and WipB. a) WipA and WipB are phosphatase effectors from *Legionella pneumophila.* b) WipA and WipB specificity were profiled using PhosPropels. c) WipA depletes pSer, pThr, and pTyr from PhosPropels at different rates. d) WipA activity on a panel of synthetic human phosphopeptides. Line shows average of n=3 biological replicates and filled area shows s.d. WipB depletes pSer, pThr, and pTyr from PhosPropels at different rates. f) WipB activity on a panel of synthetic human phosphopeptides. Line shows average of n=3 biological replicates and filled area shows s.d. g) WipA depletes phosphosites from PhosPropels with broad specificity for flanking residues, while the favorability of phosphosites as WipB substrates depends strongly on the identity of the +1 residue. h,i) WipA is a broad specificity pTyr phosphatase that depletes nearly all pTyr sites from PhosPropels over 120 min. j,k) WipB favors substrates where +1 * Pro. l) When +1 * Pro, WipB excludes acidic residues at flanking sites. (m) Alphafold3 model of the WipA dimer bound to the peptide EDAEpYAARG. (n) Alphafold3 model of the WipB dimer bound to the peptide RRALpSVASL. For panels (g), (i), and (l), z-scores were calculated by comparing positional frequencies to the 0 min timepoint using counts summed across n = 3 biological replicates. Residue-position combinations with Benjamini-Hochberg FDR-adjusted p-values < 0.0001 were considered significant and are outlined in black.

We treated PhosPropels (0.1 mg/mL) from pervanadate-treated HEK293T cells with WipA (100 nM) or WipB (100 nM) (**Fig. 3c-g**, **Supplementary Datasets 9 and 10**, **Fig. S8**, **Fig. S9**). WipA rapidly depleted pTyr sites from the library (t_1/2_ = 2.2±0.2 min), with slower activity on pSer sites (t_1/2_ = 100±13 min) and pThr sites (t_1/2_ = 19±6 min) (**Fig. 3c,g**, **Supplementary Dataset 9**). This broader activity contrasts with previous studies reporting that WipA has no activity on pSer and pThr. We validated our results using an enzyme-coupled assay to measure WipA-catalyzed phosphate release from a panel of synthetic phosphopeptides (**Fig. 3d**) and observed similarly broad activity. Given WipA’s relationship to PPP family enzymes, which typically target pSer and pThr, this broad activity is not unexpected^38^. Nonetheless, WipA’s faster kinetics toward pTyr are consistent with its proposed physiological function as a pTyr phosphatase (**Fig. 3d,h**).

We next examined sequence features associated with WipA activity on pTyr sites (**Fig. 3h, i**). At early timepoints, pTyr sites with Phe in the −1, −2, +1, and +2 positions were significantly depleted from the library, suggesting that they are good WipA substrates, while pTyr sites with acidic residues in the +1, +3, −4, and −3 positions became enriched, suggesting that they are poor substrates. However, after 2 h, <1% of pTyr sites remained, indicating that even poor substrates are eventually dephosphorylated. WipA’s broad activity raises the question of how substrate selectivity is achieved in cells. We note that WipA contains a C-terminal domain outside the phosphatase core that may direct its activity toward specific substrates or subcellular locations^35^.

WipB exhibited lower overall activity, dephosphorylating 35±5% of phosphosites within 2 h (**Fig. 3e**, **Supplementary Dataset 10**). Despite its annotation as a pSer/pThr phosphatase^37^, WipB also showed activity toward pTyr sites. Phosphosites with Pro in the +1 position became enriched in the library and persisted strongly after 2 h (**Fig. 3g**), suggesting that substrates with +1 Pro are disfavored. Phosphate release assays on a panel of five synthetic peptide substrates supported exclusion of +1 Pro, as we were unable to detect WipB-catalyzed dephosphorylation of a pSer-Pro peptide derived from aquaporin-2 (**Fig. 3f**).

To further analyze the role of +1 Pro in WipB substrate recognition, we grouped sites based on its presence or absence (**Fig. 3j, k**). Sites lacking +1 Pro were depleted from the library more rapidly, consistent with the hypothesis that these sites are preferred by WipB. Among these, pSer/pThr sites with flanking acidic residues (Asp, Glu, and pSer) persisted among the remaining phosphosites, suggesting charge-based exclusion (**Fig. 3l**). In contrast, pSer/pThr sites with aromatic (Phe, Tyr, Trp), hydrophobic (Ile, Leu, Met, Val) and polar uncharged residues at the +1, +2, and +3 positions and those with basic residues (Arg, Lys) at the +2 and +3 positions were depleted from the library. These results suggest in the absence of +1 Pro, other flanking features shape WipB specificity.

To gain insights into structural determinants of specificity, we used Alphafold3 to model dimeric WipA and WipB in complex with high-activity substrates (**Fig. 3m,n**)^39^. Consistent with an existing WipA crystal structure^35^, the top WipA model revealed a large cavity between the two subunits formed by an α-helical hairpin insertion. The pTyr residue formed extensive contacts with the active site, while flanking residues were solvent-exposed, consistent with WipA’s broad pTyr specificity. WipB, which lacks the helical insertion, formed a tighter dimer interface in which substrate contacts spanned both protomers. Glu 294 and Asp 331 are poised to interact with the substrate, potentially explaining the unfavorability of acidic substrates.

Our results reveal new insights into the substrate sequences preferred by WipA and WipB. WipA has broad activity on pTyr, pSer and pThr residues, while WipB has more restricted activity based on exclusion of +1 Pro and flanking residue context. Many preferred WipB substrates resemble motifs recognized by MAP3K, MAP4K, and NEK/ASK kinases^5^, raising the possibility that WipB may preferentially act on targets phosphorylated by these enzymes. In contrast, substrates of Pro-directed and acidophilic kinases may be resistant to WipB-catalyzed dephosphorylation. Together, our data suggest that sequence-level substrate specificity may play an important role in defining the distinct cellular functions of WipA and WipB and lay the groundwork for identifying additional physiological targets.

### PhosPropels reveal the sequences preferences of pThr lyases, a functionally and mechanistically distinct class of phosphoerasers

While some bacterial effectors subvert host signaling via phosphatase activity, others target phosphosites using mechanisms not known to occur in host cells. One example is the OspF family of pThr lyases, which catalyze β-elimination of the pThr phosphate group to generate dehydrobutyrine (Dhb) (**Fig. 4a-d**)^40^. OspF enzymes are proposed to inactivate MAP kinases (MAPKs) by targeting their activation loop pThr-Xaa-pTyr motifs, downregulating the host immune response. Previous studies have characterized OspF specificity^41,42^ but were limited in the substrate positions and amino acid substitutions examined.

**Figure 4.**
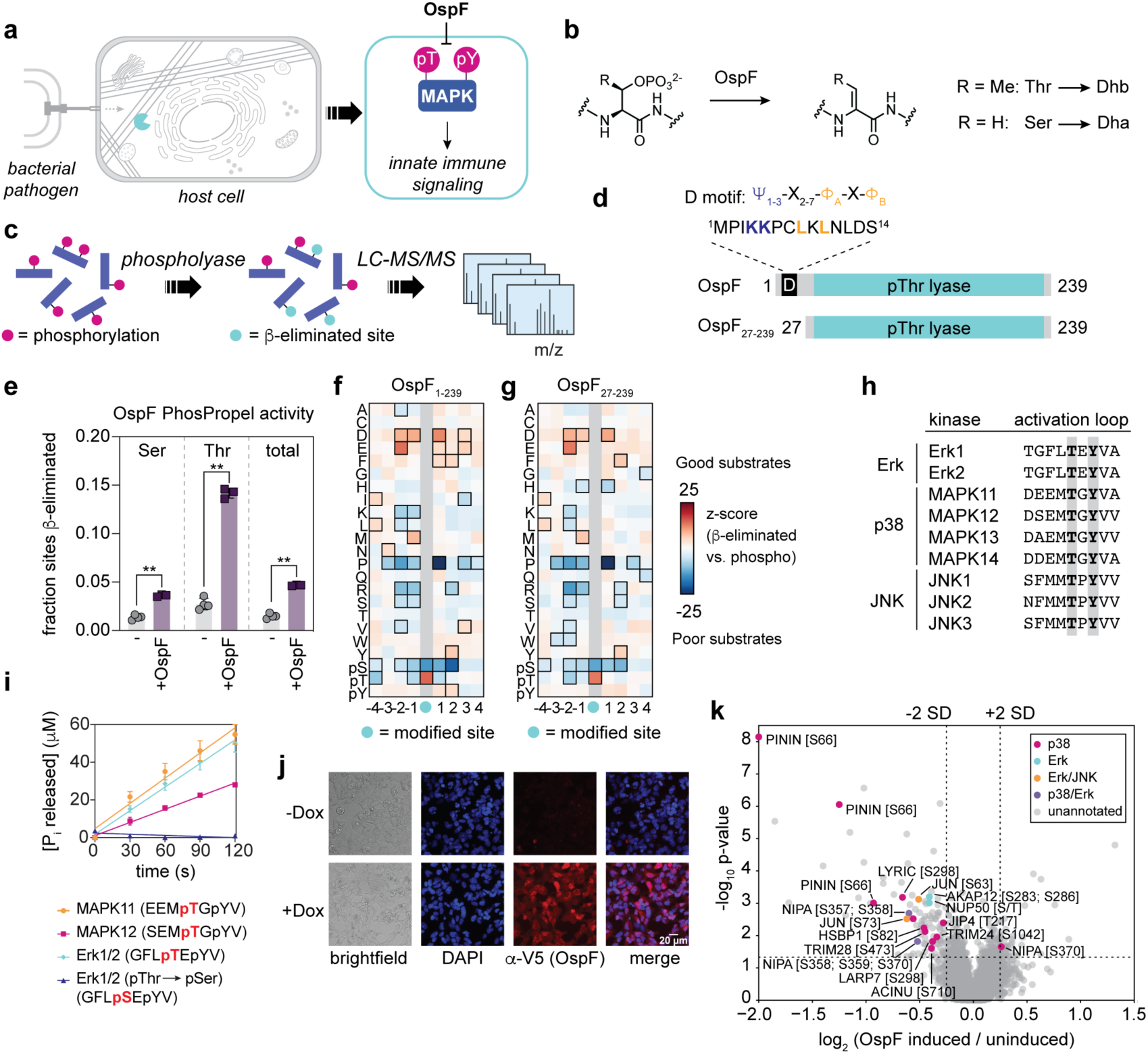
PhosPropels reveal intrinsic MAPK activation loop selectivity in OspF, a functionally distinct phosphoeraser. a) OspF is a pThr lysae effector from *Shigella flexneri*. b) OspF catalyzes β-elimination of pThr and pSer, generating Dhb or Dha. c) OspF activity on PhosPropels is detectable based on its introduction of Dha or Dhb in place of pSer or pThr, producing a −18.01 Da mass shift. d) Full-length OspF harbors a canonical D motif for MAPK docking. OspF_27-239_ has this motif deleted. e) OspF catalyzes β-elimination on PhosPropels, with higher activity on pThr residues. The fractions of sites β-eliminated in treated (purple) vs. untreated (grey) samples were compared using unpaired *t*-tests and p-values were corrected for multiple comparisons using the Holm-Šídák method. **** p < 0.0001. e) The specificity profile of full-length OspF reveals that pThr sites, sites with acidic flanking residues, and Phe, Tyr, or pTyr at +2 are favored, while +1 Pro is disfavored. g) The specificity profile of OspF_27-239_ is similar to wild-type OspF, suggesting that OspF preference for MAPK activation loop-like sequences is intrinsic to the catalytic domain. For panels (f-g), z-scores were calculated by comparing positional frequencies flanking β-eliminated sites to positional frequencies flanking phosphosites with the same sample. Counts were summed across n = 3 biological replicates. Residue-position combinations with Benjamini-Hochberg FDR-adjusted p-values < 0.0001 were considered significant and are outlined in black. h) MAPK activation loop sequences contain a pThr-Xaa-pTyr motif. In Erk, Xaa = Glu, in p38, Xaa = Gly, and in JNK, Xaa = Pro. i) OspF activity on synthetic peptides derived from MAPK activation loops. j) Doxycycline-inducible expression of OspF in HEK293T cells to probe the effect of OspF expression on the phosphoproteome. k) TMT-based quantitative proteomics reveals that OspF significantly (>2 s.d. from median phosphopeptide abundance, p<0.05) decreases the abundance of 86 phosphopeptides and increases the abundance of 23 phosphopeptides. Annotated substrates of p38 (magenta), Erk (cyan), both Erk and JNK (orange), both p38 and Erk (purple) are marked.

To comprehensively evaluate pThr lyase specificity, we used PhosPropels as substrate pools for OspF (**Fig. 4c**). After treating libraries (0.25 mg/mL) with OspF (5 μM for 24 h), we used LC-MS/MS to identify high-confidence β-elimination sites based on a −18.01 Da mass loss from Ser and Thr residues (ptmRS site localization probability >90%). OspF catalyzed β-elimination of both pSer (to form dehydroalanine, Dha) and pThr (to form dehydrobutyrine, Dhb) residues (**Fig. 4e**, **Supplementary Dataset 11**, **Fig. S10**, **Supplementary Appendix**). The frequency of Dhb sites relative to Dha sites was significantly higher than the frequency of pThr relative to pSer in untreated samples, indicating that pThr is the preferred substrate.

To define substrate preferences, we compared amino acid frequencies surrounding β-eliminated sites to the amino acid frequencies surrounding remaining pSer/pThr sites in each sample (**Fig. 4f**, **Supplementary Note 1**). OspF disfavored Pro in the +1 position, while acidic residues and Phe were most preferred. At the +2 position, aromatic residues including Phe, Tyr, and pTyr were the most preferred amino acids. N-terminal to the pSer/pThr site, acidic residues and Met at the - 1 position were preferred, while positively charged amino acids, Ser, and Pro were disfavored. An OspF homolog, SpvC, from *Salmonella enterica* had similar activity and specificity (**Extended Data Fig. 2**, **Supplementary Dataset 12**). In contrast, the pThr lyase HopAI from *Pseudomonas syringae* had little detectable activity on PhosPropels (**Extended Data Fig. 2**, **Supplementary Dataset 12**), in line with previous reports on its low activity^41^.

The sequences preferences identified using PhosPropels mirror the activation loop sequence motif of MAPKs^43,44^, which are the physiological substrates of OspF^40^. Although MAPK targeting is proposed to depend on interactions with a canonical docking peptide at the OspF N terminus (**Fig. 4d**)^45^, our experiments used short peptides, minimizing docking interactions. To test whether the catalytic domain alone is sufficient for MAPK activation loop recognition, we analyzed the specificity of an OspF variant with its docking peptide deleted (OspF_27-239_). We found that this variant retained the substrate preferences of full length OspF (**Fig. 4g**, **Supplementary Dataset 13**), supporting the conclusion that the catalytic domain harbors an intrinsic preference for MAPK activation loop sequences.

Among the three MAPK families in humans, the intervening residue between pThr and pTyr in the activation loop motif differs. In Erk1/2, this residue is Glu, while in the p38 MAPKs, it is Gly and in the JNK family, it is Pro (**Fig. 4h**)^21^. Our data suggest that Erk and p38 kinases are good substrates for OspF, while JNK is not due to the disfavored pThr-Pro motif. To test this, we measured OspF-catalyzed phosphate release from synthetic phosphopeptides derived from p38-family kinases (MAPK11 and MAPK12), Erk1/2, and a pSer variant of the Erk1/2 activation loop (**Fig. 4i**). Consistent with our phosphoproteome-derived peptide library results, the MAPK11, MAPK12, and Erk1/2 peptides were processed efficiently, while no activity was detected for the pSer peptide under the same conditions. These predictions are consistent with previous work showing that Erk1/2 and p38 are cellular OspF substrates^40,45^.

To test whether the substrate preferences that we observed are relevant in cells, we expressed OspF in HEK293T cells under a doxycycline-inducible promoter and performed tandem mass tag (TMT)-based quantitative phosphoproteomics (**Fig. 4j, k**, **Supplementary Dataset 14**). OspF induction significantly downregulated 86 phosphopeptides, while 23 phosphopeptides were upregulated. Using the PhosphoSite Plus database and relevant literature^46,47^, we queried whether downregulated phosphopeptides were downstream of Erk, p38, or JNK MAPKs. Of the significantly changed phosphosites, 16 were annotated substrates of p38, 12 were annotated substrates of Erk1/2, 2 were annotated substrates of both Erk and p38, 1 was an annotated substrate of both Erk1/2 and JNK, and 3 were annotated substrates of Erk1/2, p38, and JNK (**Fig. 4k**). Notably, none were specific to JNK, suggesting that OspF preferentially affects Erk1/2 and p38 signaling. Together, our biochemical and cellular data suggest that OspF exhibits intrinsic sequences preferences that shape its impact on the host cell phosphoproteome.

### PhosPropels enable characterization and engineering of OspF specificity

To understand the molecular basis of the observed OspF selectivity profile, we examined a crystal structure of the pThr lyase SpvC from *Salmonella enterica* bound to the peptide QYFM-pThr-E-pTyr-VA (PDB ID: 2Q8Y) (**Fig. 5a**)^45^. The peptide adopts an extended conformation, potentially explaining OspF’s poor activity on substrates with Pro at +1. The pThr phosphate group is coordinated by the residues corresponding to K102, H104, R146, R211, and R218 in OspF. The phosphate group of the pTyr residue at the +2 position interacts with K158 and K132, while the aromatic ring participates in a π-stacking interaction with F98, rationalizing OspF’s preference for +2 aromatic residues. The peptide −1 and −2 side chains are proximal to a basic pocket (R146, R211, and R218), potentially explaining the preference for acidic residues in these positions. Similarly, the +1 Glu side chain is coordinated by R213 and R218, explaining the acidic residue preference at this position.

**Figure 5.**
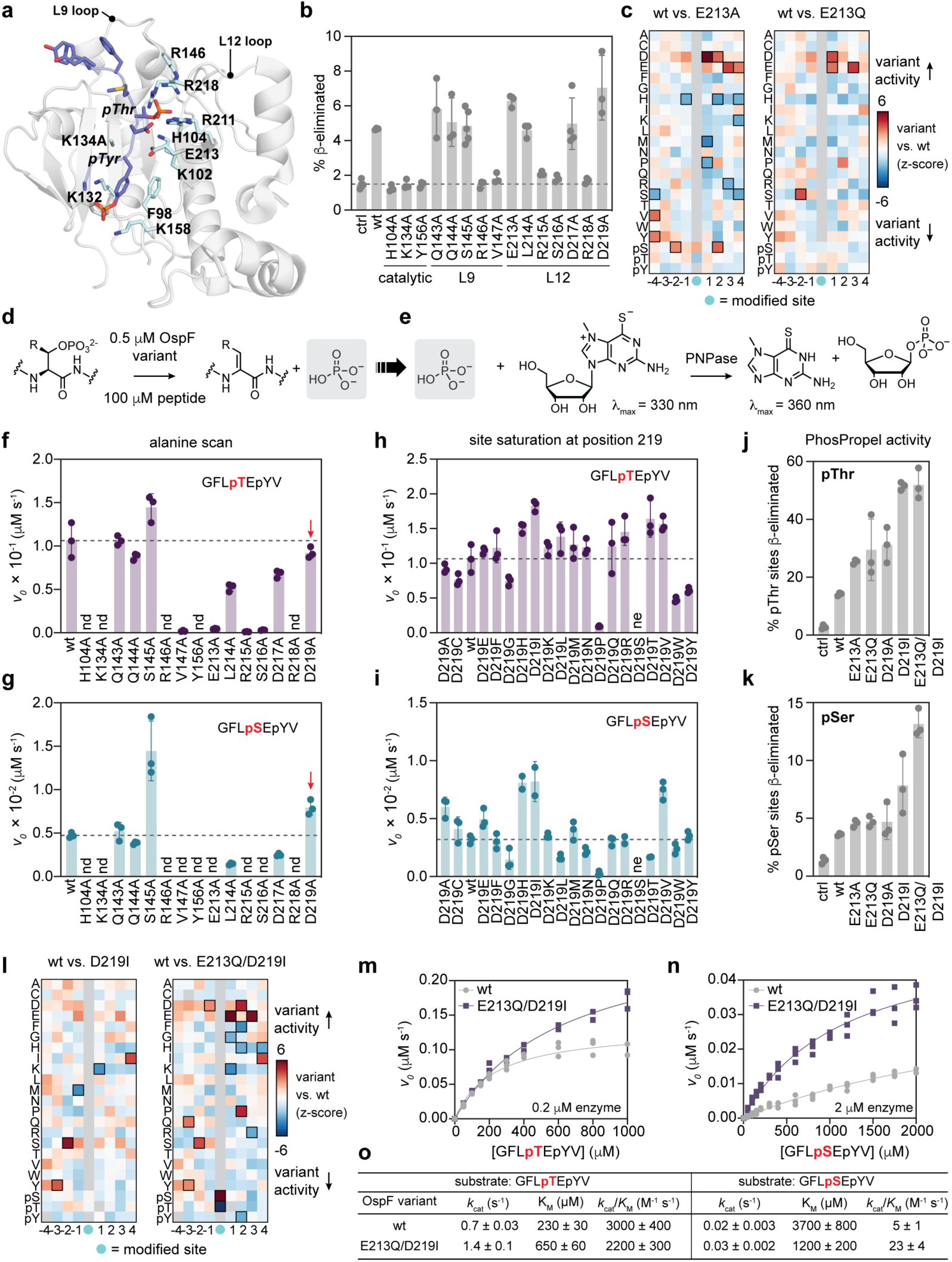
Dissecting the molecular basis of OspF sequence selectivity using PhosPropels. a) Crystal structure of OspF homolog SpvC from *Salmonella enterica* bound to the peptide QYFM-pThr-E-pTyr-VA (PDB ID: 2Q8Y). Residues are number according to OspF numbering. b) β-Elimination activity of OspF variants on PhosPropels. Dotted line indicates no-enzyme control. c) Specificity profiles of OspF-E213A and OspF-E213Q compared to wild-type. d) Screen design to measure OspF activity on GFLpTEpYV and GFLpSEpYV. e) Enzyme-coupled assay for phosphate release. f,g) OspF alanine variant activity on (f) GFLpTEpYV and (g) GFLpSEpYV. h,i) OspF-D219X activity on (h) GFLpTEpYV. and (i) GFLpSEpYV. j, k) Activity of select OspF variants on (j) pSer and (k) pThr sites in PhosPropels. l) Specificity profiles of OspF-D219I and OspF-E213Q/D219I compared to wild-type OspF. m-o) Steady-state kinetic analysis of OspF activity on (m) GFLpTEpYV and (n) GFLpSEpYV demonstrates that the variant is more efficient on the pSer peptide based on (o) an increase in *k*_cat_ and a decrease in *K*_M_. For panels (c) and (l), z-scores were calculated by comparing positional frequencies flanking β-eliminated sites in OspF variant-treated samples and wild-type OspF-treated samples. Counts were summed across n = 3 biological replicates. Residue-position combinations with Benjamini-Hochberg FDR-adjusted p-values < 0.0001 were considered significant and are outlined in black.

To probe the contribution of OspF active site residues to substrate selectivity, we performed alanine scanning mutagenesis, targeting 15 OspF residues that are predicted to be in proximity to the substrate peptide^45^. We then evaluated the activity and substrate selectivity of each variant using phosphoproteome-derived peptide libraries (**Fig. 5b**, **Supplementary Dataset 15**). Eight variants (H104A, K134A, R146A, R218A, V147A, Y156A, and S216A) had no activity above background (<3% of phosphosites β-eliminated), including three substitutions of residues that are likely directly involved in catalysis (K134A, H104A, and Y156A)^45,48,49^.

Three additional inactive variants (R215A, S216A, and V147A) substitute residues that do not directly contact the substrate, but that reside in loops that undergo conformational changes upon substrate binding (**Fig. 5a,b**)^45^. R215 and S216 residue in loop L12, which undergoes a conformational change upon substrate binding to position R218 to coordinate the pThr phosphate group. S216 hydrogen bonds with R211 to stabilize the ‘in’ conformation of the loop, while R215 hydrogen bonds with N30 in both apo and bound structures. Similarly, V147 is in loop L9, which undergoes a conformational change upon substrate binding to position R146 for coordination of the pThr phosphate. Loss of activity upon alanine substitution of these residues supports a critical role for L9 and L12 loop dynamics in OspF catalysis.

Other positions in the L9 and L12 loops were more tolerant of alanine substitution. Seven variants retained activity similar to wild-type, including Q143A, Q144A, and S145A in the L9 loop, and E213A, L214A, D217A, and D219A in the L12 loop (**Fig. 5b**). Among these residues, only E213 directly contacts the bound substrate, making hydrogen bonds with the backbone amide NH group of pTyr and the backbone carbonyl oxygen of pThr. The E213 side chain is also within hydrogen bonding distance of the +1 Glu side chain of the substrate peptide, which contains the pThr-Glu-pTyr motif characteristic of activated Erk MAPKs^21,43,44^.

To evaluate effects on specificity, we compared amino acid frequencies at positions flanking the β-eliminated sites between each active alanine variant and wild-type OspF (**Fig. 5c**, **Extended Data Fig. 3**). E213A exhibited the most substantial changes in substrate specificity (**Fig. 5c**), with a stronger preference for acidic flanking residues and a modest increase in activity toward pSer substrates. Other variants retained specificity similar to wild-type OspF, with few significant changes.

We hypothesized that substituting the E213 side chain shifts specificity by decreasing negative charge in the active site to better accommodate acidic substrates. To test this, we made the more conservative E213Q variant and analyzed its activity (**Fig. 5d**). Like E213A, E213Q has a stronger preference for acidic residues in positions flanking the β-elimination site and higher overall activity. However, the shift toward increased reactivity with pSer sites was no longer apparent, suggesting that this effect is more related to the size of the side chain in position 213 rather than its charge.

To more directly probe OspF’s selectivity for pThr over pSer, we tested the activity of the alanine variants on two synthetic peptides: the Erk2 activation loop-derived peptide GFLpTEpYV from the (**Fig. 5f**) and its pSer counterpart GFLpSEpYV (**Fig. 5g**). All mutants that were inactive on phosphoproteome-derived peptide libraries were also inactive on both of these peptides, and library-active variants had some level of activity on both peptides. While most variants had similar activity to wild-type OspF on both peptides, we found that the S145 variant had somewhat higher activity on both the pThr and pSer substrates. Notably, the D219A variant exhibited higher activity than on the pSer substrate but not the pThr substrate, suggesting that residue 219 modulates pThr vs. pSer selectivity.

To analyze the role of position 219 further, we performed saturation mutagenesis at position 219 and measured activity toward the same peptide pair (**Fig. 5h, i**). All variants except D219P retained some activity on one or both substrates. D219G severely impaired activity on the pSer substrate but not the pThr substrate, while D219H and D219I increased activity on both pSer and pThr substrates, with a more substantial increase observed for pSer. Other D219 variants had neutral or negative effects on activity. D219 is conserved in OspF, SpvC, and VirA, while it is substituted with Ala in HopA1, a low-activity homolog. These results indicate that residue 219 is a key determinant of OspF pThr selectivity.

We next analyzed selected position 219 variants in the context of PhosPropels. D219I increased total β-elimination by 2.5±0.2-fold compared to wild-type, an effect that could be attributed to increased activity on both pThr and pSer sites (**Fig. 5j,k**, **Supplementary Dataset 16**). Analysis of flanking residue preferences revealed few significant changes compared to wild-type OspF (**Fig. 5l**), indicating that the D219I variant increases β-elimination based on its overall higher activity and increased tolerance of pSer substrates.

Dha and Dhb provide versatile reactive handles that can be used for cyclization, functionalization, and post-translational mutagenesis. An efficient, broad specificity pThr/pSer lyase to install Dha/Dhb sites would be a valuable tool for these applications. Based on the broadened specificity profiles of the E213Q and D219I variants, we combined both substitutions and tested the activity of OspF-E213Q/D219I toward PhosPropels (**Fig. 5j,k,l**). The double variant β-eliminated 16±1% of phosphosites in the library, a 3.5-fold increase compared to wild-type OspF (4.7±0.1%). This level of β-elimination was also higher than either of the single variants (12±3% for D219I and 7±1% for E213Q). The increase in activity could be attributed to significant increases in β-elimination on both pSer sites and on phosphosites flanked by acidic residues (**Fig. 5l**). Kinetic analysis using GFLpTEpYV and GFLpSEpYV substrates showed that OspF-E213Q/D219I has a four-fold higher catalytic efficiency for β-elimination of the pSer substrate compared to the wild-type enzyme based on both an increase in *k*_cat_ and a decrease in *K*_M_ (**Fig. 5m,n,o**). The E213Q/D219I variant therefore combines the broadened specificity of both single mutants, providing a useful scaffold for further engineering of Dha/Dhb-generating enzymes. These results highlight how the PhosPropel approach can be deployed for protein engineering to alter enzyme specificity.

## Discussion

Enzymatic erasers of phosphorylation play central roles in cellular signaling, yet methods for profiling their sequence selectivity have remained limited in scope, throughput, or generality. To address this gap, we have developed a general and scalable strategy for specificity profiling of phosphatases and phospholyases using phosphoproteome-derived peptide libraries. A key advantage of our method is its reliance on biological samples as a regenerable source of phosphopeptides that can be readily obtained using well-established enrichment protocols. These libraries reflect the diversity of biologically relevant phosphosites and their design does not depend on prior knowledge of which motifs a particular phosphoeraser might target.

By analyzing phosphoeraser activity on phosphoproteome-derived peptide libraries, we were able to detect unexpected phosphatase sequence preferences that were overlooked in smaller scale experiments. For example, we detected substantial levels of pSer/pThr phosphatase activity in WipA and pTyr phosphatase activity in PP2Ac and WipB. Together, our results suggest that many members of the metal-dependent phosphoprotein phosphatase (PPP) enzyme family may possess broader intrinsic substrate specificity, with the higher substrate selectivity that is often observed in the cellular context arising from interactions with accessory factors and domains.

While our studies focused on phosphopeptide libraries derived from human cells, the same strategy can be extended readily to any organism or tissue from which phosphopeptides can be enriched. We showed that the composition of the library could be tuned by pervanadate pretreatment, a strategy that could be extended to any pharmacological or genetic modifier of phosphorylation signaling. Libraries could also be further tailored by additional enrichment steps (e.g., pTyr enrichment) or chemical modification (e.g., base treatment to generate dehydrated sites). More broadly, our experimental approach and statistical framework can be adapted to profile enzymes that erase or modify any post-translational modification that can be enriched for analysis by LC-MS/MS, including acetylation, methylation, and glycosylation, among many others.

The scale and complexity of phosphoproteome-derived peptide libraries enables robust statistical profiling of enzyme specificity. By comparing enzyme-treated samples to an appropriate statistical background, we systematically identified sequence features that influence substrate recognition based on their depletion or enrichment among the remaining phosphosites. This approach revealed both expected and underappreciated determinants of phosphoeraser specificity, including features that are rarely represented in synthetic phosphopeptide libraries, such as adjacent phosphosites and variable sequence lengths surrounding the phosphosite. The size and diversity of the libraries enabled us to apply targeted filters, such as restriction of the analysis to pTyr sites or exclusion of phosphosites with Pro at the +1 position, while maintaining adequate statistical power. This analytical flexibility allows specific hypotheses to be tested without requiring changes to experimental design or construction of additional libraries.

Our approach yielded new biological insights into phosphoerasers across a range of species of origin, protein folds, and enzymatic mechanisms. Using the same phosphoproteome-derived peptide libraries, we were able to characterize two classes of phosphoeraser enzymes with distinct molecular functions: phosphatases, which catalyze hydrolysis of the phosphate group to regenerate the unmodified side chain, and phospholyases, which catalyze β-elimination of the phosphate group to introduce an alkene. The throughput of our approach enabled a detailed dissection of the specificity determinants OspF, a pThr lyase. By profiling the activity and specificity of 15 alanine substitution variants of the enzyme, we identified residues that shape its substrate selectivity and catalytic efficiency. Targeted mutagenesis at key positions then allowed us to shift OspF’s toward pSer sites. These results demonstrate the power of phosphoproteome-derived peptide libraries to accelerate biological discovery by revealing principles of phosphoeraser specificity at proteome scale.

## Methods

### Key chemicals and materials

Synthetic phosphopeptides human aquaporin-2 (254-267, VELH-pSer-PQSL, cat. no. AS-61328), CK1 peptide substrate (KRRRAL-pS-VASLPGL, cat. no. AS-63797), PKC substrate 4 (KR-pThr-IRR, cat. no. AS-20291), tyrosine kinase peptide 3 (RRLIEDAE-pY-AARG, cat. no. AS-24546), and UOM9 PKC substrate (KRP-pSer-QRHG, cat. no. AS-20294) were purchased from Anaspec. Erk activation loop peptide GFL-pThr-E-pTyr-V-amide and serine variant GFL-pSer-E-pTyr-V-amide were synthesized as described previously or purchased from GenScript as the acetate salt at ≥95% purity. MAPK11 (EEM-pThr-G-pTyr-V-amide) and MAPK12 (SEM-pThr-G-pTyr-V-amide) peptides were purchased from GenScript as the acetate salt at ≥95% purity. Sequencing-grade trypsin was purchased from Promega. TiO2 beads were purchased from GL Sciences (Titansphere, 5 μm, cat. No. 5020-75000). Solid-phase extraction disks (C8 or C18, Empore) were purchased from CDS Analytical. LC-MS grade water, acetonitrile, trifluoroacetic acid (TFA), formic acid (FA), and methanol were purchased from Fisher Scientific (Optima grade). Lambda phosphatase (*λ*PP) was purchased from New England Biolabs (cat. No. P0753). Recombinant human PP2Ac (L309 deletion mutant) was purchased from Cayman Chemical (item no. 10011237). EnzChek phosphate assay kit was purchased from Thermo Fisher Scientific.

### Molecular biology and plasmid construction

*E. coli* codon-optimized genes encoding *Legionella pneumophila* WipA (RefSeq: WP_010948417.1) and WipB_1-364_ (RefSeq: WP_010946379.1), *Shigella flexneri* OspF (RefSeq: WP_010921598.1), *Salmonella typhimurium* SpvC (RefSeq: WP_001122242.1), and *Pseudomonas syringae* HopAI (RefSeq: WP_003365117.1) were purchased from Integrated DNA Technologies or Twist Bioscience. A human codon-optimized gene encoding OspF was purchased from Twist Bioscience. *E. coli* XL10 was using as the cloning host. Oligonucleotides were purchased from Integrated DNA Technologies. All plasmid sequences were confirmed via Sanger sequencing performed by Quintara Biosciences or Functional Biosciences. Plasmid maps have been deposited in the Dryad repository (DOI: 10.5061/dryad.95×69p8z0).

#### pBH4-His-Tev-OspF

The *E. coli* codon-optimized gene encoding OspF was inserted into the pBH4 vector between the BamHI and NotI restriction sites using Gibson assembly, fusing OspF with an N-terminal His tag followed by a TEV protease cleavage site. OspF mutants were generated using site-directed mutagenesis with following reaction mixture composition: forward and reverse oligonucleotides (0.5 μM each; **Table S1**), pBH4-His-Tev-OspF template (200 ng), dNTPs (0.2 mM each), MgSO_4_ (1.5 mM), KOD Hot Start DNA polymerase buffer (1×), and KOD Hot Start DNA polymerase (0.02 U/μL). The reaction mixtures were placed in a thermocycler for the following temperature cycle: 95 °C for 2 min; 25 cycles of 95 °C for 20 s, 55 °C for 10 s, 70 °C for 6 min; a final extension at 72 °C for 10 min. Reactions were then cooled and digested with DpnI (0.8 U/μl) overnight at 37 °C. *E. coli* XL10 cells were transformed with the digested products and plated on LB-agar containing carbenicillin (50 μg/mL).

#### pBH4-His-Tev-SpvC

The *E. coli* codon-optimized gene encoding SpvC was inserted into the pBH4 vector between the BamHI and NotI restriction sites using Gibson assembly, fusing SpvC with an N-terminal His tag followed by a TEV protease cleavage site.

#### pBH4-His-Tev-HopAI

The *E. coli* codon-optimized gene encoding HopAI was inserted into the pBH4 vector between the BamHI and NotI restriction sites using Gibson assembly, fusing HopAI with an N-terminal His tag followed by a TEV protease cleavage site.

#### pET28a-WipA-His

The *E. coli* codon-optimized gene encoding WipA was inserted into the pET28a vector between the NcoI and NotI restriction sites using Gibson assembly to generate a construct with a C-terminal His tag.

#### pBH4-His-Tev-WipB_1-364_

The *E. coli* codon-optimized gene encoding WipB_1-364_ was inserted into the pBH4 vector between the BamHI and NotI restriction sites using Gibson assembly to fuse WipB with an N-terminal His tag followed by a TEV protease cleavage site.

#### pcDNA5/FRT/TO-V5-OspF

The human codon-optimized gene encoding V5-OspF was inserted into the pcDNA5/FRT/TO vector (Invitrogen) between the KpnI and NotI restriction sites using Gibson assembly.

### Cell culture

HEK293T cells (ATCC #CRL-3216) and K562 cells (ATCC #CCL-243) were grown in DMEM with 10% (v/v) fetal bovine serum and 100 U/mL penicillin, 100 μg/mL streptomycin at 37°C under a 5% CO_2_ atmosphere. Jurkat E6.1 (ATCC #TIB-152) cells were grown in RPMI-1640 media with 10% fetal bovine serum and 100 U/mL penicillin, 100 μg/mL streptomycin at 37°C under 5% CO_2_ atmosphere. Flp-In TRex 293T cells (Thermo Fisher Scientific) were grown in DMEM with 10% tetracycline-free fetal bovine serum, 100 U/mL penicillin, 100 μg/mL streptomycin, and other antibiotics as appropriate. Cells were tested every six months for mycoplasma contamination using the LookOut Mycoplasma PCR Detection Kit (Sigma-Aldrich) according to the manufacturer’s instructions.

#### Construction of a stable cell line for inducible OspF expression

Flp-In TRex 293T cells were transfected with pcDNA5/FRT/TO-V5-OspF according to the manufacturer’s instructions to generate a stable cell line (Flp-In TRex 293T-V5-OspF) with V5-OspF under a doxycycline-inducible promoter.

#### Immunofluorescence

Flp-In TRex 293T-V5-OspF cells (5 × 10^4^ cells) were seeded on glass-bottom 24-well imaging dishes coated with poly-D-lysine and were grown for 24 hours at 37°C under a 5% CO_2_ atmosphere prior to induction. Media was then replaced with fresh media containing 1 μg/mL doxycycline to induce OspF expression for 18 h. Media was removed, and cells were fixed with 4% paraformaldehyde in PBS at 4°C for 45 min. Cells were then washed three times with PBS and permeabilized with cold methanol (pre-chilled at −20°C) for 5 min. Cells were washed three times with PBS and mouse anti-V5 antibody (Invitrogen #46-0705) in PBS + 3% BSA was added at a 1:2,000 dilution. Cells were incubated with the antibody for 1 h at 4°C. Cells were than washed three times with PBS and stained with DAPI (1:1,000 dilution) and goat anti-mouse-AlexaFluor 647 (1:1,000, Invitrogen #A-21236) in PBS + 3% BSA for 1 h at 4°C. Cells were washed three times with PBS prior to imaging on a Nikon Ti2-E inverted epifluorescence microscope. Raw images have been deposited in the Dryad repository (DOI: 10.5061/dryad.95×69p8z0).

### Generation of human proteome-derived peptide libraries (PhosPropels)

Human proteome-derived peptide libraries were produced following a modified version of the protocol of Jersie-Christensen et al.^17^ as described briefly below.

#### Pervanadate treatment

Pervanadate stock solution (50 mM) was made fresh prior to each use by mixing equal volumes of 100 mM hydrogen peroxide and 100 mM sodium orthovanadate. For treatment of HEK293T cells with pervanadate, media was aspirated from a 150 mm dish of cells at 90% confluency and cells were washed once with DPBS (20 mL). Serum-free DMEM (20 mL) was then added to the cells and pervanadate was added to a final concentration of 100 μM. Cells were then incubated at 37°C under 5% CO_2_ atmosphere for 1 h. Pervanadate treatment caused some cells to detach; these cells were transferred to a 50 mL centrifuge tube and harvested by centrifugation at 400 × g for 5 min. The remaining cells were treated with trypsin/EDTA (0.25%, 5 mL, Gibco) for 3 min, transferred to the same centrifuge tube, and harvested by centrifugation at 400 × g for 5 min. For Jurkat and K562 cells, cells from a 150 mm dish were collected by centrifugation at 400 × g for 5 min, washed once with DPBS, and resuspended in the appropriate serum-free media with 100 μM pervanadate. Cells were replated in a 150 mm dish and incubated at 37°C under 5% CO_2_ atmosphere for 1 h. Cells were then harvested by centrifugation at 400 × g for 5 min and serum-free media was aspirated.

#### Cell lysis

One 150 mm dish of cells at 90% confluency (HEK293T) or at a concentration of 10^6^ cells per mL (Jurkat and K562, 40 mL) was used for each replicate. The cell pellet was resuspended in 1 mL hot lysis buffer (100 mM Tris-HCl, pH 8.0, 6 M guanidine hydrochloride, 5 mM TCEP, 10 mM chloroacetamide) that was preheated to 95°C and the suspension was heated at 95°C for 10 min. Cells were then subjected to 10 cycles of probe ultrasonication (20% amplitude; 5 s on / 5 s off) to complete lysis. Insoluble material was removed by centrifugation at 20,000 × g for 10 min at 4°C.

#### Trypsin digestion

The cell lysate was diluted with 4 volumes (4 mL) of 25 mM Tris-HCl, pH 8.0 and incubated for 12-16 h at 37°C with 20 μg sequencing grade modified trypsin (Promega). The digested lysates were then acidified by adding TFA to a final concentration of 1% (v/v) to reduce the pH to <3. Insoluble material was removed by centrifugation at 5,000 × g for 5 min at 4°C. The digested peptides were desalted on a Sep-Pak C18 Plus Short Cartridge (360 mg sorbent per cartridge, Waters). The cartridge was conditioned with 100% acetonitrile (3 mL) and equilibrated with 2 × 3 mL 0.1% TFA in water. Digested peptides were then loaded onto the cartridge. The cartridge was washed with 2 × 3 mL 0.1% TFA in water. Peptides were eluted with 3 mL 40% acetonitrile/60% water/0.1% TFA followed by 3 mL 60% acetonitrile/40% water/0.1% TFA. The volume of the eluted peptides was reduced by twofold using a vacuum centrifuge. Peptide concentration was estimated by measuring absorbance at 280 nm (1 mg/mL = 1 absorbance unit, 1 cm path length) on a Nanodrop spectrophotometer.

#### Phosphopeptide enrichment

The desalted peptides were then transferred to a conical tube and its volume was doubled with the enrichment buffer (80% acetonitrile, 12% TFA). TiO_2_ bead slurry was prepared by suspending Titansphere TiO_2_ beads (GL Sciences) in bead buffer (20 mg/mL 2,5-dihydroxybenzoic acid (DHB), 80% acetonitrile, 6% TFA). The slurry was then added to the tryptic peptides in 1:2 sample:bead ratio (w/w). The sample was incubated on a rotator for 15 min and then centrifuged to collect the beads. The beads were resuspended in 50 uL of buffer A (10% acetonitrile, 6% TFA), transferred to a 1.5 mL microcentrifuge tube and centrifuged at 500 × g for 2 min. The beads were then resuspended in fresh buffer A and transferred to a homemade C8 single-layered StageTip^50^. The StageTip was centrifuged at 1,000 × g at room temperature for 2 min to remove buffer A. The StageTip was then washed with 2 × 50 μL buffer B (40% acetonitrile, 6% TFA) and 2 × 50 μL buffer C (60% acetonitrile, 6% TFA), with each wash step being carried out by centrifugation at 1,000 × g for 2 min. Phosphopeptides were eluted with 20 μL elution buffer 1 (5% ammonium hydroxide) followed by 20 μL elution buffer 2 (10% ammonium hydroxide). The eluate was concentrated in a vacuum centrifuge to near-dryness. Phosphopeptides were desalted on SOLA HRP SPE cartridges (ThermoFisher Scientific) according to the following protocol: The column was conditioned with 500 μL acetonitrile and equilibrated with 2 × 1 mL 0.1% TFA. The sample (acidified to pH < 3 with 1% TFA and diluted to 500 μL with 0.1% TFA) was loaded onto the column and column was washed with 2 × 1 mL 0.1% TFA. The desalted phosphopeptides were eluted from the column with 2 × 150 μL 80% acetonitrile with no TFA added. The sample was concentrated to near-dryness in a vacuum centrifuge, resuspended in LC-MS grade water at a concentration of 0.5-1.0 mg/mL, and stored at −20°C.

### Expression and purification of WipA and WipB

Sequences of expressed proteins are listed in **Table S2**. Chemically competent *E. coli* C43(DE3) cells were transformed with the appropriate plasmid for overexpression. Starter cultures (10 mL LB supplemented with 50 μg/mL kanamycin for WipA or 50 μg/mL carbenicillin for WipB) were inoculated with single colonies and grown overnight at 37°C with shaking at 200 rpm. Starter cultures were used to inoculate expression cultures (1 L LB supplemented with the appropriate antibiotic) in baffled flasks. Cutures were grown at 37°C with shaking at 180 rpm until they reached OD_600 nm_ of 0.4-0.6. Expression was then induced by addition of IPTG to a final concentration of 1 mM. Expression was allowed to proceed overnight at 16°C with shaking at 180 rpm. Cells were harvested by centrifugation at 4,000 × g for 20 min at 4°C and resuspended in lysis buffer (25 mM Tris-HCl, pH 7.5, 300 mM NaCl, 10 mM imidazole, 1 mM TCEP, 5% glycerol, with one Complete EDTA-Free Protease Inhibitor Tablet (Roche) per 25 mL). Cells were lysed by three passes through an Emulsiflex microfluidizer at 15,000 psi. The lysate was cleared by centrifugation at 10,000 × g for 30 min at 4°C. Clarified lysate was then added to 1 mL of Ni-NTA resin pre-equilibrated with lysis buffer. His tagged proteins were allowed to bind to the resin for 1 h on a nutator at 4°C. The resin was collected by centrifugation at 1,000 × g for 2 min at 4°C, transferred to a 15 mL column, and washed with 20 mL lysis buffer followed by 20 mL wash buffer (25 mM Tris-HCl, pH 7.5, 1 M NaCl, 10 mM imidazole, 1 mM TCEP, 5% glycerol). Protein was then eluted with 2-3 × 5 mL elution buffer (25 mM Tris-HCl, pH 7.5, 150 mM NaCl, 600 mM imidazole, 1 mM TCEP, 5% glycerol). For WipB, TEV protease (1:50 molar ratio) was added prior to dialysis again dialysis buffer (25 mM Tris-HCl, pH 7.5, 150 mM NaCl, 1 mM TCEP, 5% glycerol) overnight. WipA was dialyzed against the same buffer in the absence of TEV protease. After dialysis, WipB was passed over a Ni-NTA column to removed uncleaved protein and protease. Purified proteins were aliquoted, flash frozen in liquid nitrogen, and stored at −80°C until futher use. Purity was assessed by SDS-PAGE and ESI-TOF MS analysis (**Figure S11**).

### Expression and purification of OspF, SpvC, HopAI, and OspF variants

Sequences of expressed proteins are listed in **Table S2.** Chemically competent *E. coli* BL21(DE3) cells were transformed with the appropriate plasmid for overexpression. Starter cultures (10 mL Luria-Bertani (LB) media, 50 μg/mL carbenicillin) were inoculated with single colonies and grown overnight at 37°C with shaking at 200 rpm. Starter cultures were used to inoculate expression cultures (1 L LB, 50 μg/ml carbenicillin) in baffled flasks. Cultures were shaken at 180 rpm at 37°C until they reached OD_600_ ∼0.6. Expression was then induced with 1 mM IPTG (isopropyl-β-D-thiogalactopyranoside). The expression was carried out overnight at 16°C with shaking at 180 rpm. Cell pellets were harvested by centrifugation and resuspended in 40 mL lysis buffer (25 mM Tris pH 8.0, 200 mM NaCl, 5 mM imidazole, pH 8.0, 300 μM TCEP). Lysis was achieved by three passes through an Emulsiflex microfluidizer at 15000 psi. The lysate was centrifuged at 10,000 × g for 30 min at 4°C. Batch binding with HisPur Ni-NTA resin (1 mL, ThermoFisher Scientific) was allowed for 1 h at 4°C on a nutator. Afterwards, the resin was collected by centrifugation at 1,000 × g for 2 min at 4°C, transferred to a 15 mL column, and washed with 20 mL lysis buffer and 20 mL wash buffer (25 mM Tris pH 8.0, 300 mM NaCl, 20 mM imidazole, pH 8.0, 300 μM TCEP). Protein was eluted with 10 mL elution buffer (25 mM Tris pH 8.0, 150 mM NaCl, 250 mM imidazole, pH 8.0, 300 μM TCEP) and dialyzed overnight against 2 L dialysis buffer (25 mM Tris pH 8.0, 150 mM NaCl, 300 μM TCEP). Dialyzed protein was concentrated in an Amicon Ultra-4 centrifugal filter (10,000 NMWL) (Millipore) prior to purification by size exclusion chromatography on Superdex 75 Increase 16/60 column (Cytiva) using size exclusion buffer (25 mM Tris pH 8.0, 150 mM NaCl, 300 μM TCEP). Fractions containing pure recombinant protein were pooled, concentrated, and flash frozen in liquid nitrogen. Following the purification, the protein purity was analyzed by SDS-PAGE and ESI-TOF MS (**Figure S12**). The protein concentration was determined by absorbance at 280 nm, with extinction coefficients determined by ProtParam.

### Treatment of PhosPropels with phosphoeraser enzymes

Specific conditions for reactions with each enzyme are described below. After quenching, reactions were desalted using double-layer C18 StageTips made in house according to the protocol of Rappsilber et al.^50^. Buffers and samples were passed through the StageTip by centrifugation at 1,000 × g for 1 min. The tip was conditioned with 50 μL of acetonitrile and equilibrated with 2 × 50 μL 0.1% TFA. The sample was then loaded. The tip was then washed with 2 × 50 μL 0.1% TFA, 1 × 50 μL 5% methanol, 0.1% TFA, and 1 × 50 μL 2% acetonitrile, 0.1% FA. Desalted peptides were eluted with 2 × 30 μL 80% acetonitrile, 0.1% FA. Samples were concentrated to near-dryness in a vacuum centrifuge and then resuspended in 13 μL 2% acetonitrile, 0.1% FA. The concentration was estimated by measuring the sample absorbance at 280 nm on a Nanodrop spectrophotomer (1 mg/mL = 1 absorbance unit with a 1 cm path length).

#### *λ* phosphatase (*λ*PP)

*λ*PP was purchased from New England Biolabs (cat. No. P0753). PhosPropels (0.1-0.2 mg/mL) were treated with *λ*PP (30 units) in 50 mM HEPES, pH 7.5, 100 mM NaCl, 2 mM DTT, 1 mM MnCl_2_ at 30°C in a total volume of 50 μL. The reaction was quenched after 5 min, 30 min, or 180 min by addition of 10 μL of 1%TFA.

#### PTP1B_1-321_

PTP1B_1-321_ was purified as described previously^51^. PhosPropels (0.1-0.2 mg/mL) were treated with PTP1B_1-321_ (20 nM) in 10 mM Tris-HCl, pH 7.5, 25 mM NaCl, 1 mM EDTA, 1 mM DTT at 37°C in a total volume of 50 μL. The reaction was quenched after 5 min, 30 min, or 60 min by addition of 10 μL of 1%TFA.

#### PP2Ac

Recombinant human PP2Ac (L309 deletion mutant) was purchased from Cayman Chemical (item no. 10011237). PhosPropels (0.1-0.2 mg/mL) were treated with PP2Ac (250 nM) in 40 mM Tris-HCl, pH 7.5, 34 mM MgCl_2_, 4 mM EDTA, 2 mM DTT, 0.05 mg/mL BSA at 30°C in a total volume of 50 μL. The reaction was quenched after 5 min, 30 min, or 180 min by addition of 10 μL of 1%TFA.

#### WipA

PhosPropels (0.1-0.2 mg/mL) were treated with WipA (100 nM) in 25 mM Tris-HCl, pH 7.5, 150 mM NaCl, 1 mM DTT, 1 mM MnCl_2_ at 37°C in a total volume of 50 μL. The reaction was quenched after 5 min, 30 min, or 120 min by addition of 10 μL of 1%TFA.

#### WipB

PhosPropels (0.1-0.2 mg/mL) were treated with WipB (100 nM) in 25 mM Tris-HCl, pH 7.5, 150 mM NaCl, 1 mM DTT, 1 mM MnCl_2_ at 37°C. The reaction was quenched after 5 min, 30 min, or 120 min by addition of 10 μL of 1%TFA.

### LC-MS/MS data collection for PhosPropel samples

LC-MS/MS data were collected using an UltiMate 3000 RSLCnano liquid chromatography system in line with an Orbitrap Exploris 480 hybrid quadrupole-Orbitrap mass spectrometer (ThermoFisher Scientific). Peptides (0.5-1 μg dissolved in 2% acetonitrile, 0.1% FA, 5 μL total injection volume) were injected onto an Acclaim PepMap RSLC column (75 µm × 50 cm, 2 µm particle size, 100 Å pore size, ThermoFisher Scientific) over 15 min in 97% mobile phase A (0.1% FA) and 3% mobile phase B (0.1% FA, 80% acetonitrile) at 0.3 μL/min. The column was eluted using a linear gradient from 3% mobile phase B to 50% mobile phase B over 120 min at 0.3 μL/min. The eluate was electrosprayed through a nanospray emitter tip by applying 2000 V through the ion source’s DirectJunction adapter. Full MS scans were performed over a range of 350-1,200 m/z at a resolution of 60,000 at 200 m/z. The AGC target was set to 300% and the maximum injection time was set to ‘auto’. The top 20 most abundant precursors with a charge state of 2-6 were selected for MS/MS analysis using an isolation window of 1.4 m/z and a precursor intensity threshold of 5 × 10^3^. A 20 s dynamic exclusion window with a precursor mass tolerance of ± 10 ppm was applied. MS/MS scans used HCD fragmentation with a normalized collision energy of 30%, a resolution of 15,000 and a fixed first mass of 110 m/z. The AGC target was set to ‘standard’ and the maximum injection time was set to 22 ms.

### LC-MS/MS data analysis for PhosPropels

Thermo RAW files were searched against the human SwissProt database (downloaded 01/24/2020)^52^ using the SEQUEST^53^ algorithm in Proteome Discoverer 2.4 (ThermoFisher Scientific). The precursor mass tolerance was set to 10 ppm and the fragment mass tolerance was set to 0.02 Da. Search parameters included the following modifications: carbamidomethylation at Cys (+57.021 Da, static), oxidation at Met (+15.995 Da, dynamic), acetylation at protein N termini (+42.011 Da, dynamic), Met loss at protein N termini (−131.040 Da, dynamic), Met loss+acetylation at protein N termini (−89.030 Da, dynamic), phosphorylation at Ser, Thr, and Tyr (+79.966 Da, dynamic), and dehydration at Ser and Thr (−18.011 Da, dynamic). Up to two missed cleavages were allowed. PSM validation was performed with the Percolator^54^ node of Proteome Discoverer 2.4 at a false discovery rate of 1%. Localization of modification sites was scored using the IMP-ptmRS^55^ node with PhosphoRS set to false. Raw data, peak lists, and results have been deposited in the ProteomeXchange repository under the accession numbers listed in **Table S3**. Results are also provided in Microsoft Excel format as **Supplementary Datasets S1-S16** as described in **Table S3.**

### Analysis of PhosPropel data for phosphoeraser specificity profiling

Z-scores for each amino acid in each position were calculated using custom Python scripts available in the Zenodo repository (DOI: 10.5281/zenodo.16785439). Z-scores compared the position-specific frequencies of each amino acid in each position surrounding the phosphosite to frequencies in an appropriate experimentally measured background set. For each position and amino acid, z-scores were calculated as described below. Z-scores were assumed to follow a standard normal distribution under the null hypothesis, which is appropriate given the large number of observations contributing to each frequency estimate. Z-scores are plotted directly in the heatmaps shown for each specificity profile, colored according to their magnitude and direction. Two-tailed p-values were computed from the z-scores using the standard normal distribution. Multiple hypothesis testing correction was performed using the Benjamini-Hochberg method to control the false discovery rate at a = 0.0001. Position-amino acid combinations that had statistically significant frequency differences between the sample and the control are outlined in black in the heatmaps. The statistical framework for this analysis is described in detail in **Supplementary Note 1**.

#### PhosPropel positional composition

To generate the z-score heatmaps shown in **Fig. 1D**, phosphosites identified with ptmRS localization scores >90 were aligned and the positional frequencies of each amino acid in each position were calculated. The global (position-independent) frequency of each amino acid (including pSer, pThr, and pTyr) observed in each sample was then calculated. Z-scores were then calculated according to the equation

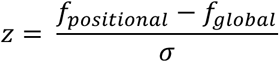

where f_positional_ is the frequency of each amino acid in each position surrounding the phosphosite, f_global_ is the global frequency, and σ is the population standard deviation,

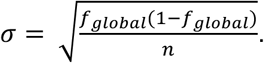

#### PhosPropel phosphatase specificity profiles

To generate the z-score heatmaps shown for phosphatase specificity profiles, phosphosites identified with ptmRS localization scores >90 were aligned and the positional frequencies of each amino acid in each position were calculated. Z-scores were then calculated by comparing positional frequencies at each time point to positional frequencies at the 0 min time point according to the equation

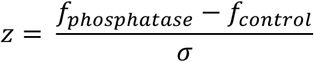

where f_phosphatase_ is the frequency of each amino acid in each position surrounding the phosphosite following phosphatase treatment for the specified time, f_control_ is the frequency of each amino acid in each position at the 0 min time point, and σ is the population standard deviation,

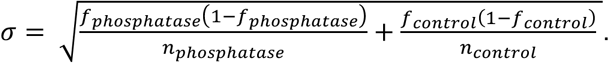

#### PhosPropel phospholyase specificity profiles

To generate the z-score heatmaps shown for phospholyase specificity profiles, β-elimination sites identified with ptmRS localization scores >90 were aligned and the positional frequencies of each amino acid in each position surrounding the β-elimination site were calculated. Separately, pSer and pThr sites with ptmRS localization scores >90 were aligned and the positional frequencies of each amino acid in each position surrounding the phosphosite were calculated. Only pSer and pThr sites were used because β-elimination can only occur at pSer and pThr sites. Z-scores were then calculated by comparing positional frequencies for β-elimination sites versus phosphosites according to the equation

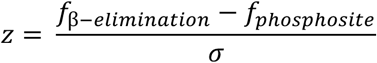

where f_β-elimination_ is the frequency of each amino acid in each position surrounding β-elimination sites, f_phosphosite_ is the frequency of each amino acid surrounding phosphosites, and σ is the population standard deviation,

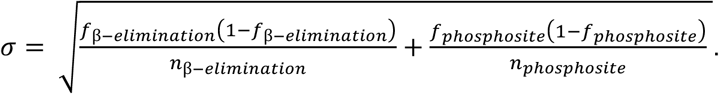

#### Comparison between OspF variant and wild-type OspF specificity profiles

To generate the z-score heatmaps shown for comparison of specificity profiles between OspF variants and wild-type OspF, β-elimination sites identified with ptmRS localization scores >90 in the OspF variant-treated samples were aligned and the positional frequencies of each amino acid in each position surrounding the β-elimination site were calculated. Separately, β-elimination sites identified with ptmRS localization scores >90 in the wild-type OspF-treated samples were aligned and the positional frequencies of each amino acid in each position surrounding the β-elimination site were calculated. Z-scores were then calculated by comparing positional frequencies for β-elimination sites in the two samples according to the equation

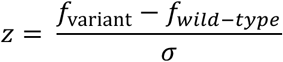

where f_variant_ is the frequency of each amino acid in each position surrounding β-elimination sites in the wild-type OspF-treated sample, f_wild-type_ is the frequency of each amino acid in each position surrounding β-elimination sites in the wild-type OspF-treated sample, and σ is the population standard deviation,

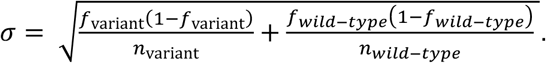

### Scoring of Phospropel phosphosites with kinase PSSMs

Phosphopeptides identified in each Phospropel replicate were scored using position-specific scoring matrices (PSSMs) for Ser/Thr kinases (for pSer/pThr sites) or for Tyr kinases (for pTyr sites) using custom code that is available in the Zenodo repository under DOI: 10.5281/zenodo.16785439. Each site was scored against all kinases by summing the PSSM values reported in ref. ^5,6^ for each position-amino acid combination in a window four residues upstream and four residues downstream of the site. Kinases with a score ≥2 standard deviations above the mean score for a given sequence were considered as candidate writers of the phosphosite and are included in the dataset. For each replicate, the number of times a given kinase passed this threshold (i.e., the number of potential substrates per replicate) was computed. The average number of substrates per sample was then calculated. The log_2_(average number of substrates per replicate) was plotted on the human kinome tree as node size using Coral^56^. For comparison of the average number of potential kinase substrates in untreated and pervanadate-treated libraries, *P* values were calculated using a two-tailed, unpaired *t* test in Prism 10 (GraphPad). *P* values were corrected for multiple comparisons using the Holm-Sidak method available in the Prism 10 software.

### Generation of Alphafold3 models of substrate-bound WipA and WipB

Structural models of WipA and WipB were generated using Alphafold3^39^ accessed at https://alphafoldserver.com/. For WipA, the model included two molecules of WipA, two molecules of the peptide EDAEpYAARG (a validated WipA substrate), and two Mn^2+^ ions. For WipB, the model included two molecules of full-length WipB, two molecules of the peptide RRALpSVASL, and two Mn^2+^ ions. Models were ranked based on their predicted LDDT (pLDDT) confidence scores, and the highest-confidence structure was used for analysis. Models were rendered in PyMOL and have been deposited in the Dryad repository (DOI: 10.5061/dryad.95×69p8z0).

### Preparation of TMT phosphoproteomics samples

HEK293T OspF Flp-In cells were grown in 150 mm culture plates in the presence or absence of 1 μg/mL doxycycline for 24 hours. Four biological replicates were performed for each expression condition. Cells were washed with PBS and treated with trypsin/EDTA (0.25%, 5 mL, Gibco) for 3 min, and harvested by centrifugation at 400 × g for 5 min. Proteomic lysis buffer (1 mL per 150 mm culture dish, 100 mM Tris-HCl, pH 8, 6 M guanidine-HCl, 7.5 mM TCEP, 5 mM chloroacetamide) was heated to 95°C then added to cell pellets. Cells in lysis buffer were heated for an additional 10 minutes and then sonicated (10 cycles at 20% amplitude; 5 s on / 5 s off). Lysates were diluted with 25 mM Tris, pH 8.0 such that guanidine-HCl was below 1 M then digested using 20 μg sequencing-grade trypsin (Promega) at 37°C overnight.

Peptide samples were desalted using the Sep-Pak Plus C18 cartridge (Waters) desalting procedure described in **Generation of human proteome-derived peptide libraries (PhosPropels)** above. Dried, desalted peptides were resuspended in 100 mM HEPES, pH 8 and further pH adjusted using 1M HEPES until an approximate pH of 8 was reached. Peptide concentration was assessed using the Pierce Quantitative Colorimetric Peptide Assay Kit (Thermo Fisher Scientific). 400 μg of peptide from each sample was labeled with 800 μg TMT10plex isobaric labeling reagent (Thermo Fisher Scientific) resuspended in acetonitrile. After incubation at room temperature for 1hr, reactions were quenched using 0.27% hydroxylamine for 15 minutes. Samples were then pooled and dried in a vacuum centrifuge. The dried pooled sample was again desalted using the standard SepPak desalting procedure.

The pooled TMT labeled sample was then enriched for phosphopeptides as described in **Generation of human proteome-derived peptide libraries (PhosPropels)** above. Dried, TMT-labeled phosphopeptide samples were desalted using standard SOLA C18 HRP SPE cartridge (Thermo Fisher Scientific) procedure.

### LC-MS/MS data collection for TMT phosphoproteomics samples

TMT phosphoproteomics samples were analyzed on a Thermo Scientific Ultimate 3000 RSLCnano coupled to a Thermo Scientific Orbitrap Lumos Tribrid. Sample was resuspended 0.1% FA / 5% acetonitrile, and 1 μg of sample was loaded onto a Thermo Fisher Scientific EASY-spray column (500 mm length × 75 μm inner diameter, 2 μm particle size, spherical fully porous ultrapure silica). Peptides were eluted using mobile phase A (0.1% FA) and mobile phase B (80% acetonitrile / 0.1% FA) with the gradient profile: 0 min: 2% B, 3 min: 5% B, 140 min: 25% B, 206min: 62.5% B, 216 min: 95% B. The Lumos MS was operated in positive mode using a 1.8 kV spray voltage and 300°C ion transfer tube temperature. MS1 spectra were collected at 60,000 resolution for a m/z range of 400 – 1,400. MS2 spectra were collected in a 1 s total cycle time using a 0.7 m/z isolation width, 36% normalized HCD collisional energy, 30,000 resolution and a mass range of 110 – 2,000 m/z. A maximum injection time of 50 ms and normalized AGC target of 250% was used for both MS1 and MS2 spectra collection. Dynamic exclusion was set to 30s.

### TMT phosphoproteomics data analysis

LC-MS/MS data was processed using the Sequest HT^53^ algorithm in Proteome Discoverer 2.4 (ThermoFisher Scientific). Data were searched using the human SwissProt FASTA database (downloaded 1/24/2020)^52^ with a precursor mass tolerance of 10 ppm and a fragment mass tolerance of 0.6 Da. Static modifications of TMT6plex (+229.163 Da, peptide N-terminal and K) and carbamidomethyl (+57.021 Da, C), as well as dynamic modifications acetyl (+42.011 Da, protein N-terminal), Met-loss (−131.040 Da, M), Met-loss+Acetyl (−89.030 Da, M), oxidation (+15.995 Da, M), and phosphorylation (+79.966 Da, S,T,Y). High confidence peptides were identified with a 0.01 target FDR and medium confidence peptides were identified with a 0.05 target FDR. Quantification of peptide reporter ions was performed in Proteome Discoverer 2.4 (ThermoFisher Scientific) and was normalized based on total peptide amount per channel and p-values were calculated using an ANOVA test based on individual peptides.

### Phosphate release assays

Kinetic analysis of phosphate release catalyzed by phosphoerasers was performed using the EnzChek Phosphate Assay Kit (ThermoFisher Scientific) following manufacturer’s protocol, with changes in buffer composition according to the requirements of the enzyme under study (described in detail below). Reactions were performed in 100 μL total volume. Assays were conducted in UV-Star UV-Transparent Microplates (Greiner) and absorbance at 360 nm was monitored for 30 min with a Tecan Infinite M200 Pro microplate reader.

#### Single substrate concentration rate measurements

Assays were performed using peptides (AQP2, CK1, PKC, TKP, and UOM9, Erk, Erk (pThr→pSer), MAPK11, or MAPK12) at a final concentration of 100 μM. PTP_1-321_ was assayed at 20 nM in 1× reaction buffer provided in the EnzChek kit. PP2Ac was assayed at 4 mU/mL in 1× reaction buffer provided in the EnzChek kit. WipA was assayed at 200 nM in 1× reaction buffer supplemented with 1 mM MnCl_2_. WipB was assayed at 300 nM in 1× reaction buffer supplemented with 1 mM MnCl_2_. OspF was assayed at 200 nM using a custom reaction buffer (300 mM Tris-HCl, pH 7.5, 1 mM MgCl_2_).

#### OspF variant activity screen

Peptides GFLpTEpYV and GFLpSEpYV were used at 100 μM final concentration. For alanine scan, OspF was assayed at 0.5 μM for the pThr susbtrate and 2 μM for the pSer substrate. OspF-D219X variants were assayed at a final concentration of 0.5 μM in 300 mM Tris-HCl, pH 7.5, 1 mM MgCl_2_ for both substrates.

#### Michaelis-Menten kinetics of OspF and OspF-D213Q/E219I

To determine steady-state kinetic parameters of OspF and OspF-E213Q/D219I, substrate concentration was varied between 0-1000 μM for GFLpTEpYV and between 0-2000 μM for GFLpSEpYV. Phosphopeptides were added to a reaction mixture containing 300 mM Tris-HCl, pH 7.5, 1 mM MgCl_2_, 0.2 mM 2-amino-6-mercapto-7-methylpurine riboside (MESG), and 0.1 U purine nucleoside phosphorylase (PNP). The reactions were started by addition of OspF or OspF-E213Q/D219I (0.2 μM for pThr-containing substrate, 2 μM for pSer-containing substrate) and immediately placed in the plate reader. Initial rates for each substrate concentration were determined by linear fitting. Replicate initial rates were plotted versus substrate concentration and fit to the Michaelis-Menten equation to determine *k*_cat_ and *K*_M_ using non-linear regression in GraphPad Prism 10.

## Supporting information

Supplementary Dataset 1 - HEK293T untreated phosphopeptide library

Supplementary Dataset 2 - HEK293T pervanadate-treated phosphopeptide library

Supplementary Dataset 3 - Jurkat pervanadate-treated phosphopeptide library

Supplementary Dataset 4 - K562 pervanadate-treated phosphopeptide library

Supplementary Dataset 5 - Average kinase substrate counts based on PSSM scores

Supplementary Dataset 6 - HEK293T pervanadate-treated phosphopeptide library treated with lambda phosphatase

Supplementary Dataset 7 - HEK293T pervandate-treated phosphopeptide library treated with PTP1B catalytic domain

Supplementary Dataset 8 - HEK293T pervanadate-treated phosphopeptide library treated with PP2Ac

Supplementary Dataset 9 - HEK293T pervanadate-treated library treated with WipA.

Supplementary Dataset 10 - HEK293T pervanadate-treated library treated with WipB

Supplementary Dataset 11 - HEK293T pervanadate-treated library treated with OspF

Supplementary Dataset 12 - HEK293T pervanadate-treated library treated with SpvC or HopAI

Supplementary Dataset 13 - HEK293T pervanadate-treated library treated with OspF 27-239

Supplementary Dataset 14 - OspF TMT results

Supplementary Dataset 15 - OspF alanine variant activity on HEK293T phosphopeptide library

Supplementary Dataset 16 - Additional OspF variant activity on HEK293T phosphopeptide library

## Data availability

Tandem mass spectrometry data have been deposited in the ProteomeXchange repository under the accession numbers PXD067199, PXD067202, PXD067205, PXD067206, PXD067209, and PXD067267 and are publicly available. Plasmid maps, uncropped microscopy images, and Alphafold3 models have been deposited in the Dryad repository with the DOI: 10.5061/dryad.95×69p8z0 and are publicly available.

## Code availability

The code used to perform analyses and generate figures is available at https://github.com/aweeks8/phospropel and has been archived on Zenodo with DOI: 10.5281/zenodo.16785439. An interactive notebook with examples workflows is available via Google Colab: https://colab.research.google.com/drive/1z0EJLxZw_3K2Fey708cgrAeFc4LOzC47?usp=sharing.

## Acknowledgements

We thank S. Coyle, D. Sashital, T. Galateo, members of the Weeks lab, and Ajinkya Kokandakar of the UW-Madison Statistical Consulting Group for helpful discussions. We thank Grzegorz Sabat and Dr. Greg Barret-Wilt of the UW-Madison Biotechnology Center Mass Spectrometry/Proteomics Facility for assistance with TMT phosphoproteomics data acquisition. This work was supported in part by startup funds from the University of Wisconsin-Madison Department of Biochemistry and by an NIH Director’s New Innovator Award (DP2GM149548) to A.M.W. L.E.M. was supported in part by the UW-Madison Biotechnology Training Program under grant number NIH 5 T32 GM135066 and by William H. Peterson and Dr. Steven Babcock Agricultural Chemistry Graduate Fellowships from the University of Wisconsin-Madison Department of Biochemistry.

## Extended Data Figures and Legends

**Extended Data Figure 1.**
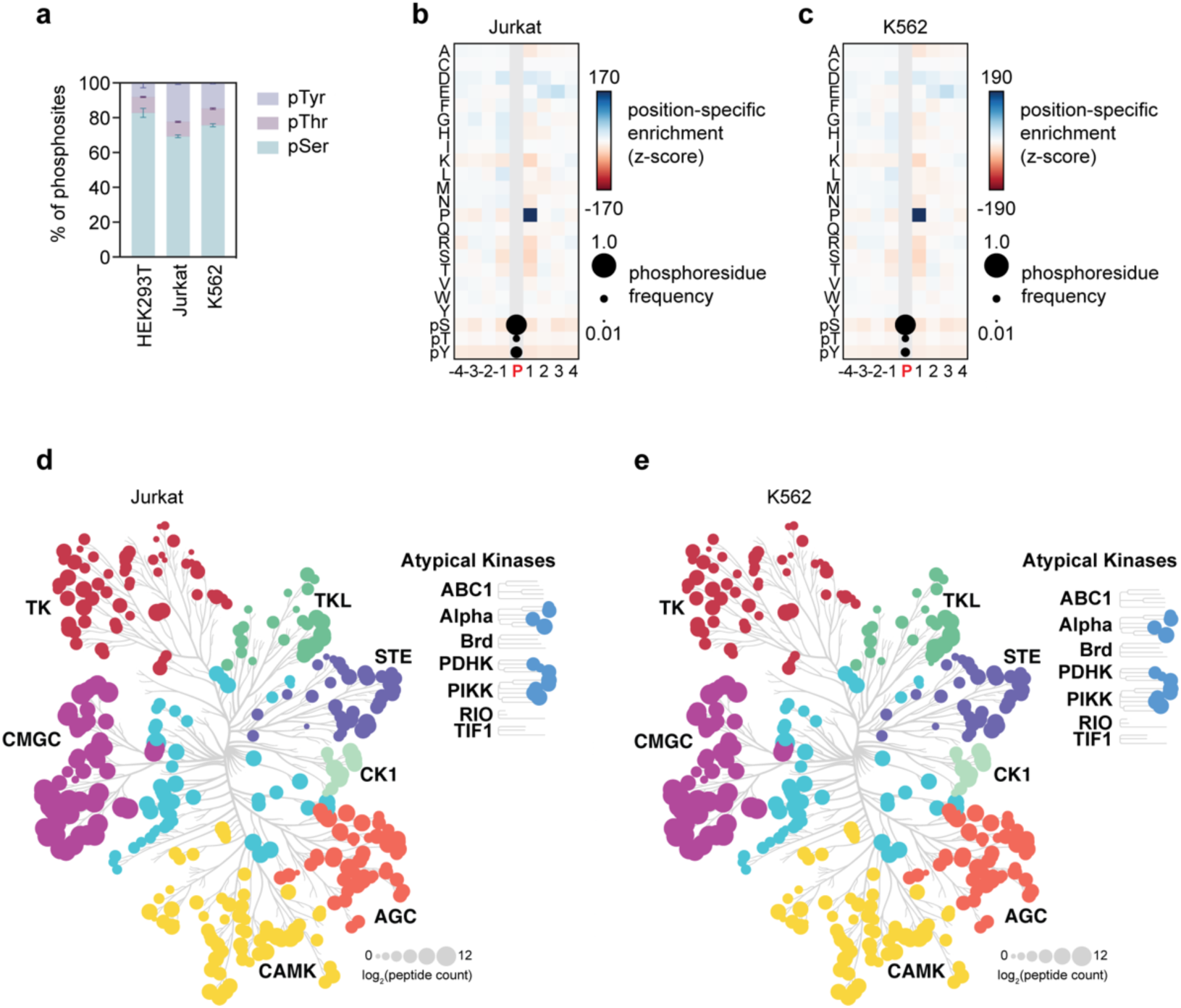
Phosphoproteome-derived peptide libraries (PhosPropels) from pervanadate-treated Jurkat and K562 cells. a) Distribution of pSer, pThr, and pTyr sites in PhosPropels from pervanadate-treated HEK293T, Jurkat, and K562 cells. b) Z-score heatmap showing the distribution of all 20 proteinogenic amino acids, pSer, pThr, and pTyr in positions flanking phosphsites in the Jurakt PhosPropel. c) Z-score heatmap showing the distribution of all 20 proteinogenic amino acids, pSer, pThr, and pTyr in positions flanking phosphsites in the Jurakt PhosPropel. d,e) Distrubution of high-scoring kinase PSSM scores among phosphopeptides in the Jurkat (d) and K562 (e) PhosPropels. Node size represents the average log_2_ count of sequences that score at least 2 standard deviations above the kinase-specific PSSM mean.

**Extended Data Figure 2.**
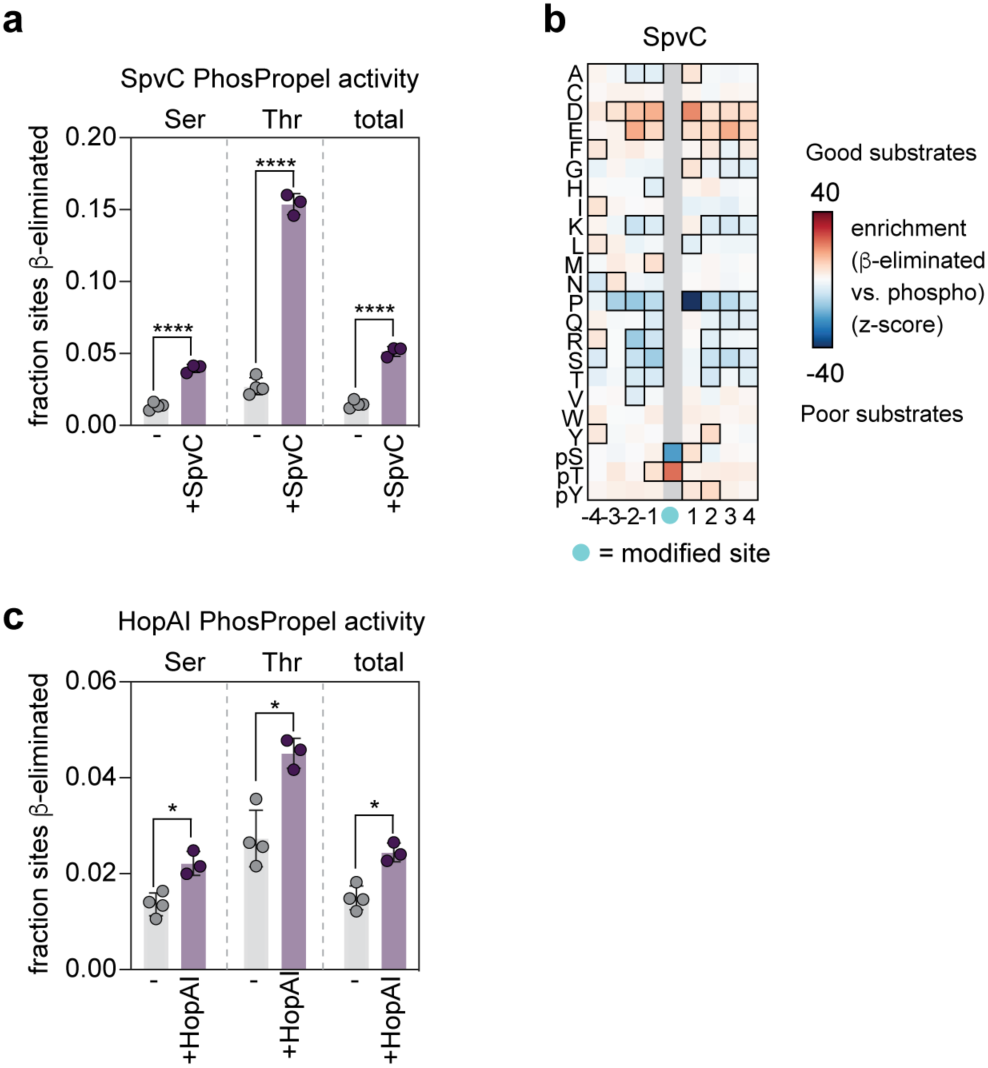
Characterization of the pThr lyases SpvC and HopAI using PhosPropels. a) SpvC catalyzes β-elimination on PhosPropels with a similar specificity profile to OspF. The fractions of sites β-eliminated in treated (purple) vs. untreated (grey) samples were compared using unpaired *t*-tests and p-values were corrected for multiple comparisons using the Holm-Šídák method. **** p < 0.0001. b) The specificity profile of SpvC reveals that pThr sites, sites with acidic flanking residues, and Phe, Tyr, or pTyr at +2 are favored, while +1 Pro is disfavored. c) HopAI catalyzes β-elimination on PhosPropels with lower activity than OspF or SpvC. The fractions of sites β-eliminated in treated (purple) vs. untreated (grey) samples were compared using unpaired *t*-tests and p-values were corrected for multiple comparisons using the Holm-Šídák method. * p < 0.05.

**Extended Data Figure 3.**
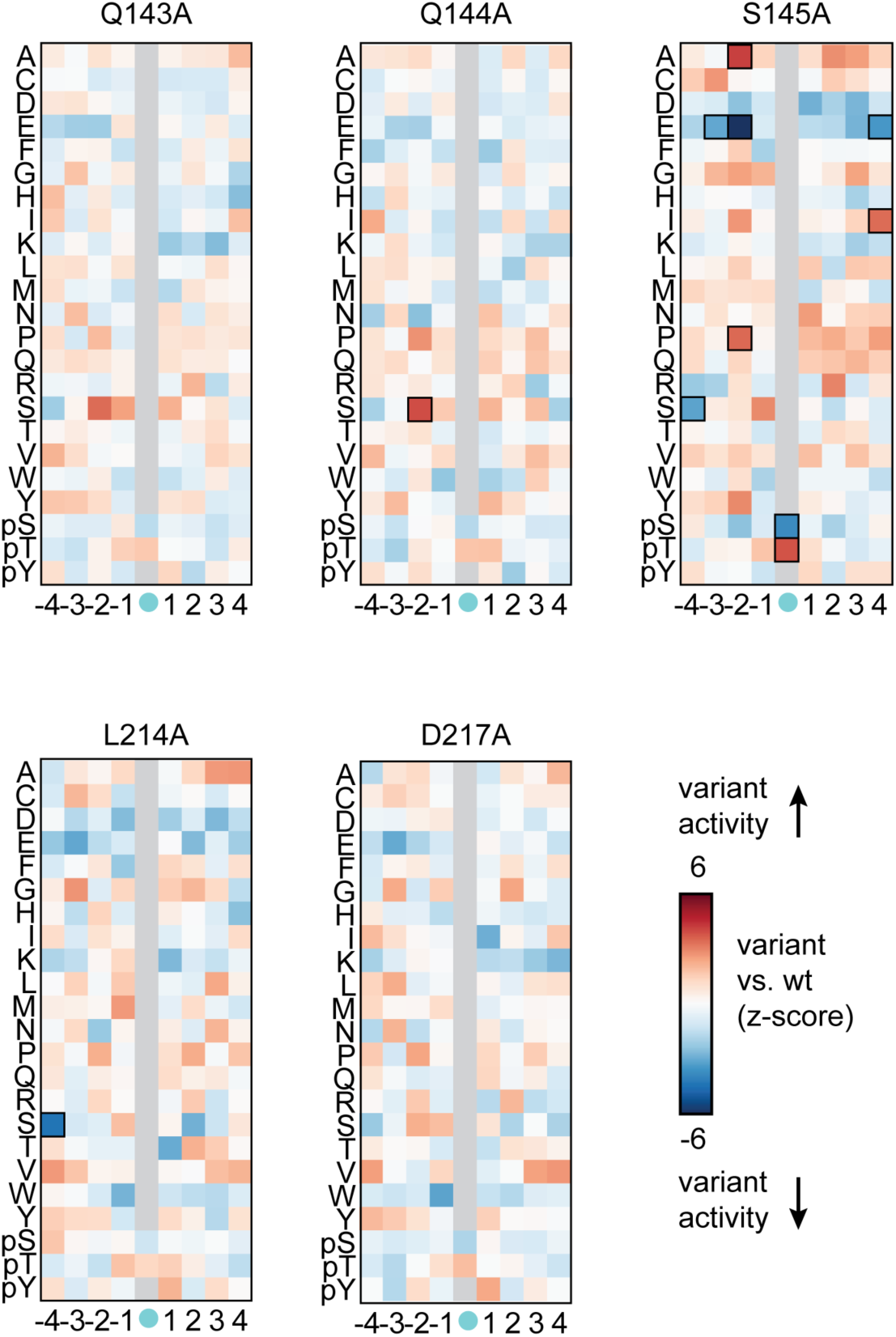
Specificity profiles of Osp alanine variants compared to wild-type OspF. Z-scores were calculated by comparing positional frequencies flanking β-eliminated sites in OspF variant-treated samples and wild-type OspF-treated samples. Counts were summed across n = 3 biological replicates. Residue-position combinations with Benjamini-Hochberg FDR-adjusted p-values < 0.0001 were considered significant and are outlined in black.

