## Supporting Tables and Figures for "Phosphoproteome-derived peptide libraries for deep specificity profiling of phosphatases and phospholyases"

**Table S1. Sequences of primers used in site-directed mutagenesis.**

| Variant | Forward primer sequence | Reverse primer sequence |
|---|---|---|
| H104A | GTTGGGGACAAATTTGCAATTAGCATCGCT CGC | GCGAGCGATGCTAATTGCAAATTTGTCCCCAAC |
| K134A | CCTATTGATAAGTGGGCAATTACGGACATG AAT | ATTCATGTCCGTAATTGCCCACTTATCAATAGG |
| Q143A | ATGAATCGCGTCTCCGCACAATCTCGCGTT GGG | CCCAACGCGAGATTGTGCGGAGACGCGATTCA T |
| Q144A | AATCGCGTCTCCCAAGCATCTCGCGTTGG GATT | AATCCCAACGCGAGATGCTTGGGAGACGCGATT |
| S145A | CGCGTCTCCCAACAAGCACGCGTTGGGAT TGGT | ACCAATCCCAACGCGTGCTTGTTGGGAGACGC G |
| S145X | TGAATCGCGTCTCCCAACAANNKCGCGTT GGGA | TCCCAACGCGMNNTTGTTGGGAGACGCGATTC A |
| R146A | GTCTCCCAACAATCTGCAGTTGGGATTGGT GCT | AGCACCAATCCCAACTGCAGATTGTTGGGAGAC |
| V147A | TCCCAACAATCTCGCGCAGGGATTGGTGC TCAG | CTGAGCACCAATCCCTGCGCGAGATTGTTGGGA |
| Y156A | GCTCAGTTTACTTTGGCAGTAAAGTCCGAC CAA | TTGGTCGGACTTTACTGCCAAAGTAAACTGAGC |
| E213A | GTTTCATATCGTAATGCATTACGCTCAGAT CGT | ACGATCTGAGCGTAATGCATTACGATATGAAAC |
| E213Q | AAGTACGTTTCATATCGTAATCAATTACGC TCAGATCGTGATG | CATCACGATCTGAGCGTAATTGATTACGATATGA AACGTACTT |
| L214A | TCATATCGTAATGAAGCACGCTCAGATCGT GAT | ATCACGATCTGAGCGTGCTTCATTACGATATGA |
| R215A | TATCGTAATGAATTAGCATCAGATCGTGAT GGC | GCCATCACGATCTGATGCTAATTCATTACGATA |
| S216A | CGTAATGAATTACGCGCAGATCGTGATGG CTCC | GGAGCCATCACGATCTGCGCGTAATTCATTACG |
| D217A | AATGAATTACGCTCAGCACGTGATGGCTC CGAA | TTCGGAGCCATCACGTGCTGAGCGTAATTCATT |
| R218A | GAATTACGCTCAGATGCAGATGGCTCCGA ACGT | ACGTTCGGAGCCATCTGCATCTGAGCGTAATTC |
| D219A | TTACGCTCAGATCGTGCAGGCTCCGAACG TCAG | CTGACGTTCGGAGCCTGCACGATCTGAGCGTAA |
| D219X | TTACGCTCAGATCGTNNKGGCTCCGAACG TCAG | CTGACGTTCGGAGCCMNNACGATCTGAGCGTA A |

**Table 2. Sequences of proteins expressed in the study.**

| Protein name | Protein sequence |
|---|---|
| WipA | MPKRLINKNIDIYNYPNEFEDNLGSISLGDLHGNAIKLIHFLFRHKIIKFKTEII<br>NFHEAYQQFVTIYEQYDDMVQEYLEIRTLLQLIQIKITNAQQRILDIEQKLSL<br>ATDHQKEFSQSLLQLKKPIEANLQMAEKSKAGLEEKLSGLKTRLPSCIERF<br>NKFMTQIEINDIKTLIRLLGDEVADRGSCDYFTLRILDFLYQNQIAIKIILSNHG<br>YEFIHAYEKLVVGQPFKPKGYIGDIQIKSFWGLQLLLEQSVITEEELRSLVER<br>AYKPTLKIIDYSLSEDGITLYSHAPIRFDSIRMAASQLGVTYNDSTKEALAETI<br>DQLNAQLQIYMKNNMLHLLFENNEINDPTNMTDEERNASPLIYLVWNRWN<br>ESKEVENARPGKYNGYFVTYVHGHDPFQSPLTYVYNLDTLCGKYSRVGE<br>EEQINKAFQFLTENRHTNVDKTASESLLRNISRYKVLDSDEYTLKHKIPKTS<br>LELATDILDCKIKESLIKLSLLGKPKAVSGSLSVDQNISIPNQASLGKGHHHH<br>HH |
| WipB | MGHHHHHHDYDIPTTENLYFQGSMKTQRIIHPNIDIRKFPEVNTDFSMTDIS<br>MGDLHANALLFLNILVRQGIIAISPENYAKFAEIYTLPELQADYWGTEAPVFS<br>AENKQERLEEIKKQYNALIAQIKIINTKKLIRLIGDELVDRGVIDYFILKLLQAL<br>YDQGADFEILLSNHGIEFVEACELFKENGNKLVAKRLGNIQHGNSFHALQE<br>AIAAGAISNEEVLNIYHQVYKKHLKIISYSLDPDANEIKVFSHAGIGLNHIRGL<br>ARKFKVPYSEESAVDLAKTIDAINKKFAEKASSGEIHTLYTHDMMYRGYAG<br>EHLNSTDEVVAATVWGREYGDLIRTSKKFKITFIHGHDSYDPEKVEHVTLN<br>NQLGQFQNNVGDLYLYATNG |
| OspF | MGHHHHHHDYDIPTTENLYFQGSMPIKKPCLKLNLDSLNVVKSEIPQMLSA<br>NERLKNNFNILYNQIRQYPAYYFKVASNVPTYSDICQFFSVMYQGFQIVNH<br>SGDVFIHACRENPQSKGDFVGDKFHISIAREQVPLAFQILSGLLFSEDSPID<br>KWKITDMNRVSQQSRVGIGAQFTLYVKSDQECSQYSALLLHKIRQFIMCLE<br>SNLLRSKIAPGEYPASDVRPEDWKYVSYRNELRSDRDGSERQEQMLREE<br>PFYRLMIE |
| SpvC | MGHHHHHHDYDIPTTENLYFQGSMPINRPNLNLNIPPLNIVAAYDGAEIPST<br>NKHLKNNFNSLHNQMRKMPVSHFKEALDVPDYSGMRQSGFFAMSQGFQ<br>LNNHGYDVFIHARRESPQSQGKFAGDKFHISVLRDMVPQAFQALSGLLFS<br>EDSPVDKWKVTDMEKVVQQARVSLGAQFTLYIKPDQENSQYSASFLHKT<br>RQFIECLESRLSENGVISGQCPESDVHPENWKYLSYRNELRSGRDGGEM<br>QRQALREEPFYRLMTE |
| HopAI | MGHHHHHHDYDIPTTENLYFQGSMNPIQSRFSSVQELRRSNVDIPALKAN<br>GQLEVDGKRYEIRAADDGTISVLRPEQQSKAKSFFKGASQLIGGSSQRAQI<br>AQALNEKVASARTVLHQSAMTGGRLDTLERGESSSATTAIKPTAKQAAQS<br>TFNSFHEWAKQAEAMRNPSRMDIYKIYKQDAPHSHPMSDEQQEEFLHTL<br>KALNGKNGIEVRTQDHDSVRNKKDRNLDKYIAESPDAKRFFYRIIPKHERR<br>EDKNQGRLTIGVQPQYATQLTRAMATLIGKESAITHGKVIGPACHGQMTDS<br>AVLYINGDVAKAEKLGEKLKQMSGIPLDAFVEHTPLSMQSLSKGLSYAESIL<br>GDTRGHGMSRAEVISDALRMDGMPFLARLKLSLSANGYDPDNPALRNTK |
| $OspF_{25-239}$ | MGHHHHHHDYDIPTTENLYFQGSMLSANERLKNNFNILYNQIRQYPAYYF<br>KVASNVPTYSDICQFFSVMYQGFQIVNHSGDVFIHACRENPQSKGDFVGD<br>KFHISIAREQVPLAFQILSGLLFSEDSPIDKWKITDMNRVSQQSRVGIGAQF<br>TLYVKSDQECSQYSALLLHKIRQFIMCLESNLLRSKIAPGEYPASDVRPED<br>WKYVSYRNELRSDRDGSERQEQMLREEPFYRLMIE |

**Table S3.** List of ProteomeXchange accession numbers and experimental information.

| Experiment # | experiment description | Relevant to | Supplementary Dataset # | ProteomeXchange # |
|---|---|---|---|---|
| Exp1 | Phosphoproteome-derived peptide libraries from untreated HEK293T cells | Fig. 1 | 1 | PXD067199 |
| Exp2 | Phosphoproteome-derived peptide libraries from pervanadate-treated HEK293T cells | Fig. 1 | 2 | PXD067199 |
| Exp3 | Phosphoproteome-derived peptide libraries from pervanadate-treated K562 cells | Fig. 1 | 4 | PXD067199 |
| Exp4 | Phosphoproteome-derived peptide libraries from pervanadate-treated Jurkat cells | Fig. 1 | 3 | PXD067199 |
| Exp5 | Timecourse of $\lambda$PP treatment of Phospropel from pervanadate-treated HEK293T cells | Fig. 1 | 6 | PXD067199 |
| Exp6 | Timecourse of PTP1B$_{1\text{-}321}$ treatment of Phospropel from pervanadate-treated HEK293T cells | Fig. 2 | **7** | PXD067202 |
| Exp7 | Timecourse of PP2Ac treatment of Phospropel from pervanadate-treated HEK293T cells | Fig. 2 | 8 | PXD067202 |
| Exp8 | Timecourse of WipA treatment of Phospropel from pervanadate-treated HEK293T cells | Fig. 3 | 9 | PXD067205 |
| Exp9 | Timecourse of WipB treatment of Phospropel from pervanadate-treated HEK293T cells | Fig. 3 | 10 | PXD067205 |
| Exp10 | OspF treatment of Phospropel from pervanadate-treated HEK293T cells | Fig. 4 | 11 | PXD067206 |
| Exp11 | SpvC treatment of Phospropel from pervanadate-treated HEK293T cells | Fig. 4 | 12 | PXD040046 |
| Exp12 | HopAI treatment of Phospropel from pervanadate-treated HEK293T cells | Fig. 4 | 12 | PXD040046 |
| Exp13 | OspF$_{25\text{-}239}$ treatment of Phospropel from pervanadate-treated HEK293T cells | Fig. 4 | 13 | PXD040046 |
| Exp14 | Inducible OspF TMT experiment | Fig. 4 | 14 | PXD067267 |
| Exp15 | Treatment of phosphoproteome-derived peptide libraries with OspF Ala variants | Fig. 5 | 15 | PXD067209 |

| | | | | |
|---|---|---|---|---|
| Exp 16 | Treatment of phosphoproteome-derived peptide libraries with additional OspF variants | Fig. 5 | 16 | PXD067209 |

**Figure S1. Distribution of ptmRS site localization scores in PhosPropels from untreated HEK293T cells.** Scores were computed using the IMP-ptmRS node in Proteome Discoverer 2.4 (ThermoFisher Scientific). Only sites with scores >90 were included in further analysis.

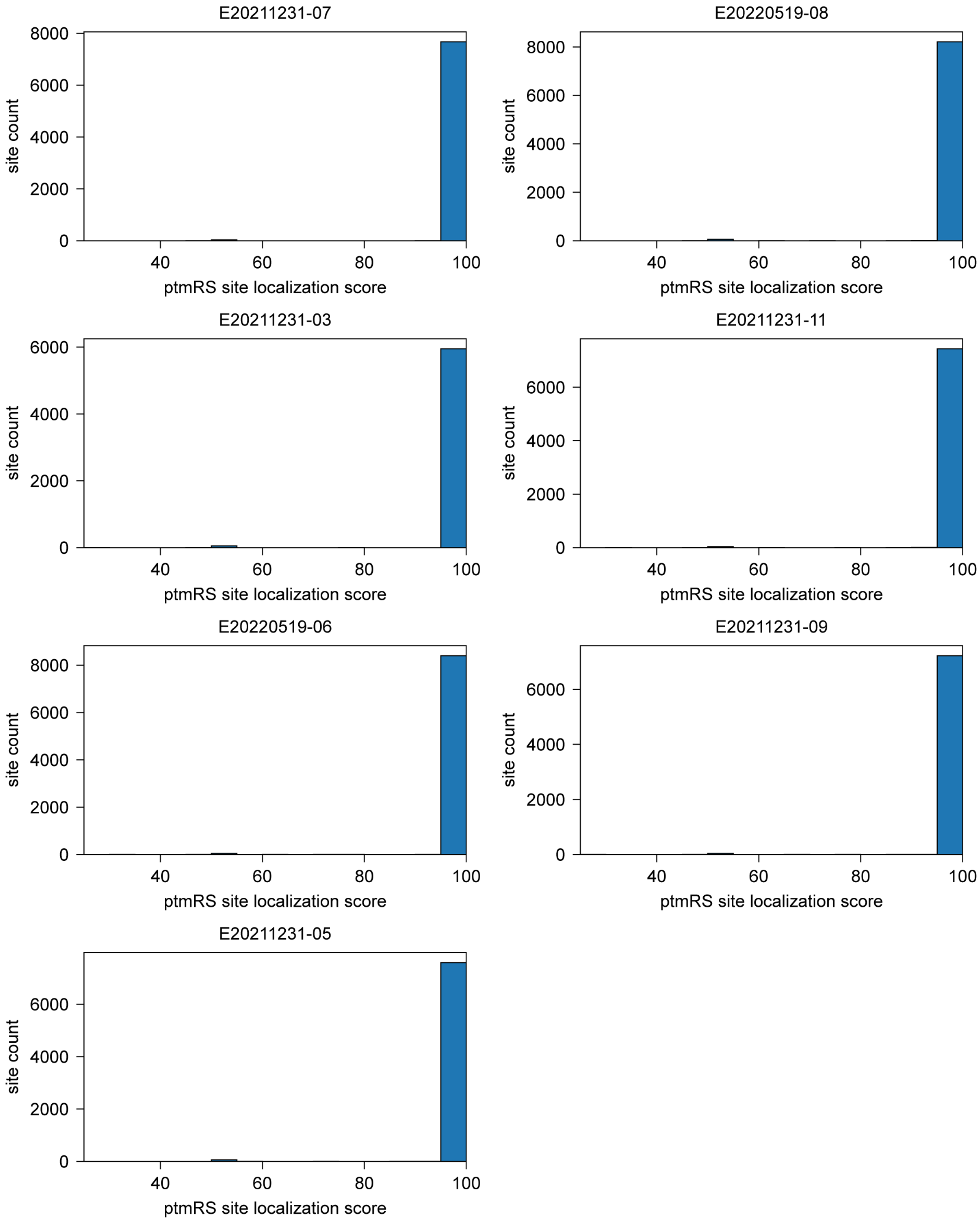

**Figure S2. Distribution of ptmRS site localization scores in PhosPropels from pervanadate-treated HEK293T cells.** Scores were computed using the IMP-ptmRS node in Proteome Discoverer 2.4 (ThermoFisher Scientific). Only sites with scores >90 were included in the analysis.

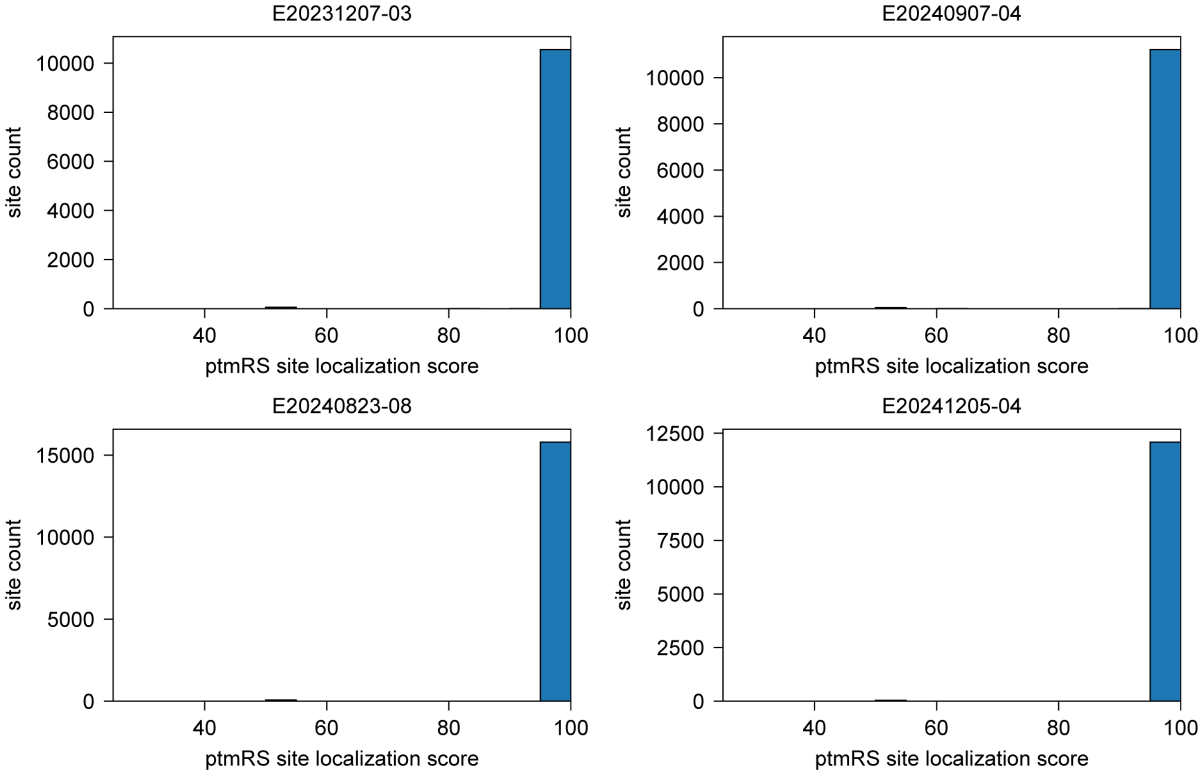

**Figure S3. Distribution of ptmRS site localization scores in PhosPropels from pervanadate-treated Jurkat cells.** Scores were computed using the IMP-ptmRS node in Proteome Discoverer 2.4 (ThermoFisher Scientific). Only sites with scores >90 were included in the analysis.

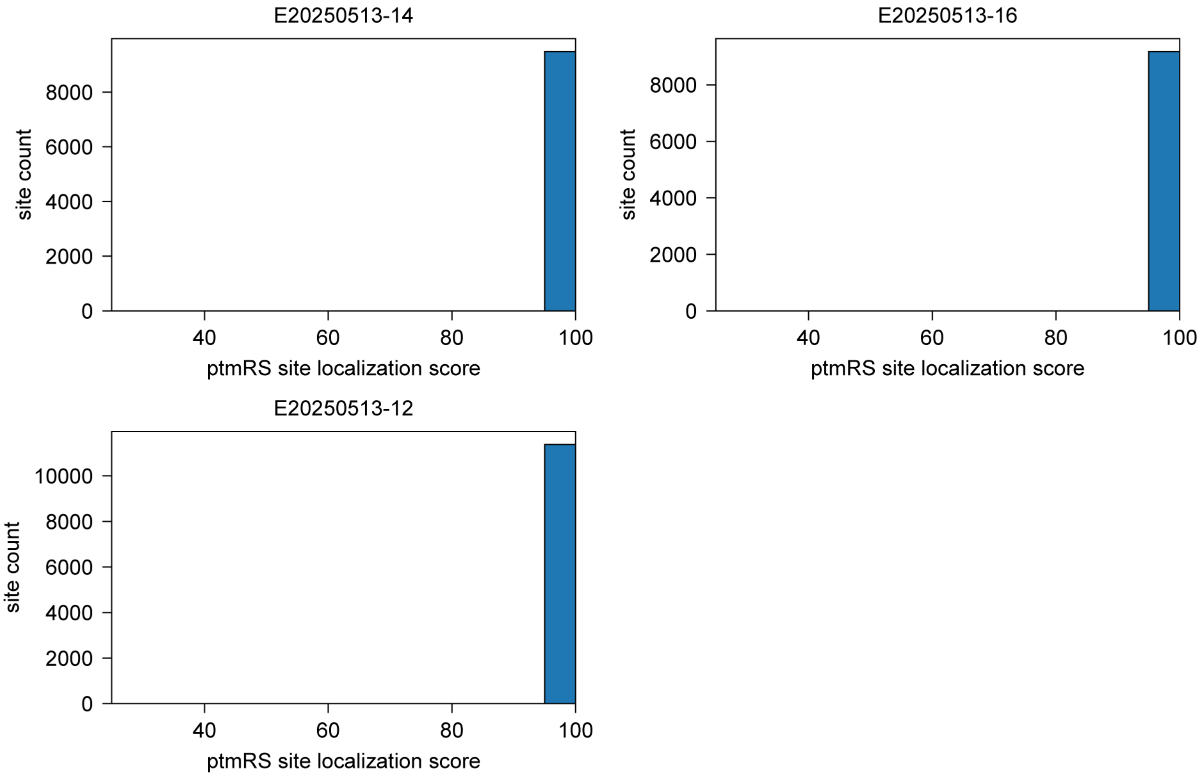

**Figure S4. Distribution of ptmRS site localization scores in PhosPropels from pervanadate-treated K562 cells.** Scores were computed using the IMP-ptmRS node in Proteome Discoverer 2.4 (ThermoFisher Scientific). Only sites with scores >90 were included in the analysis.

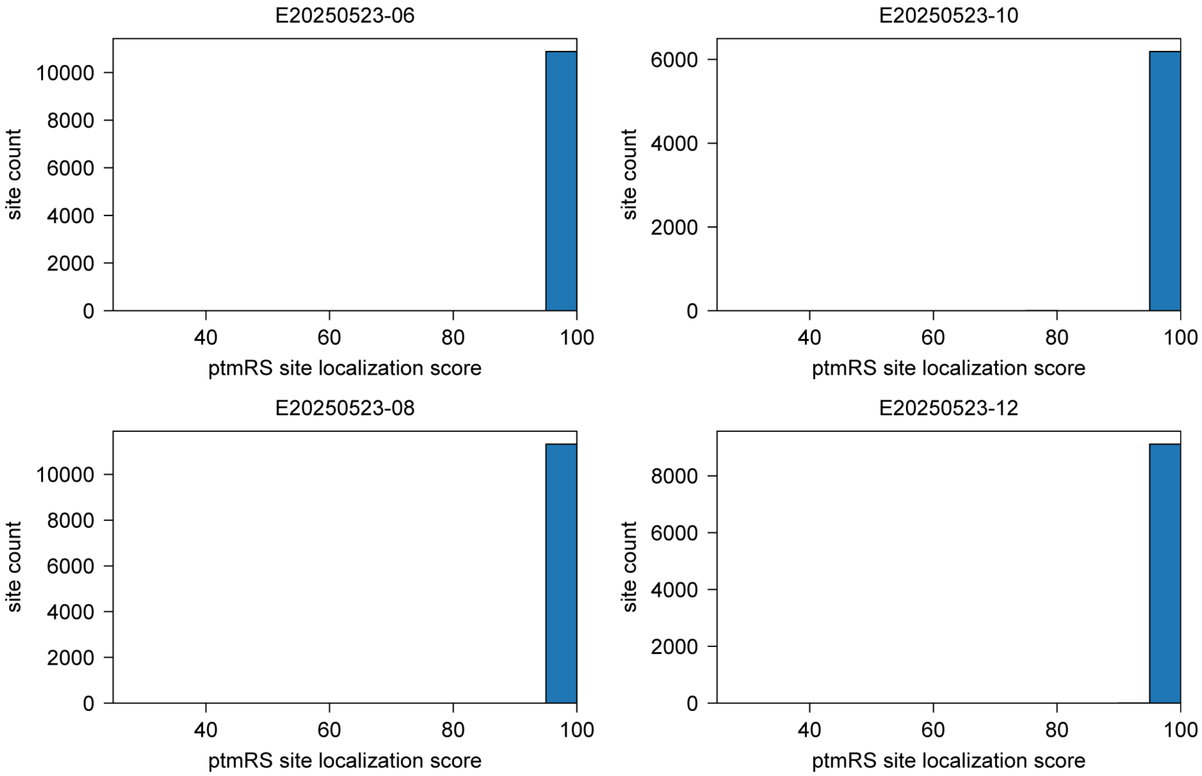

**Figure S5. Representation of pSer, pThr, and pTyr residues in positions flanking a central phosphosite.** Bars show mean ± s.d for untreated (grey) and pervanadate-treated (purple) PhosPropels from HEK293T cells.

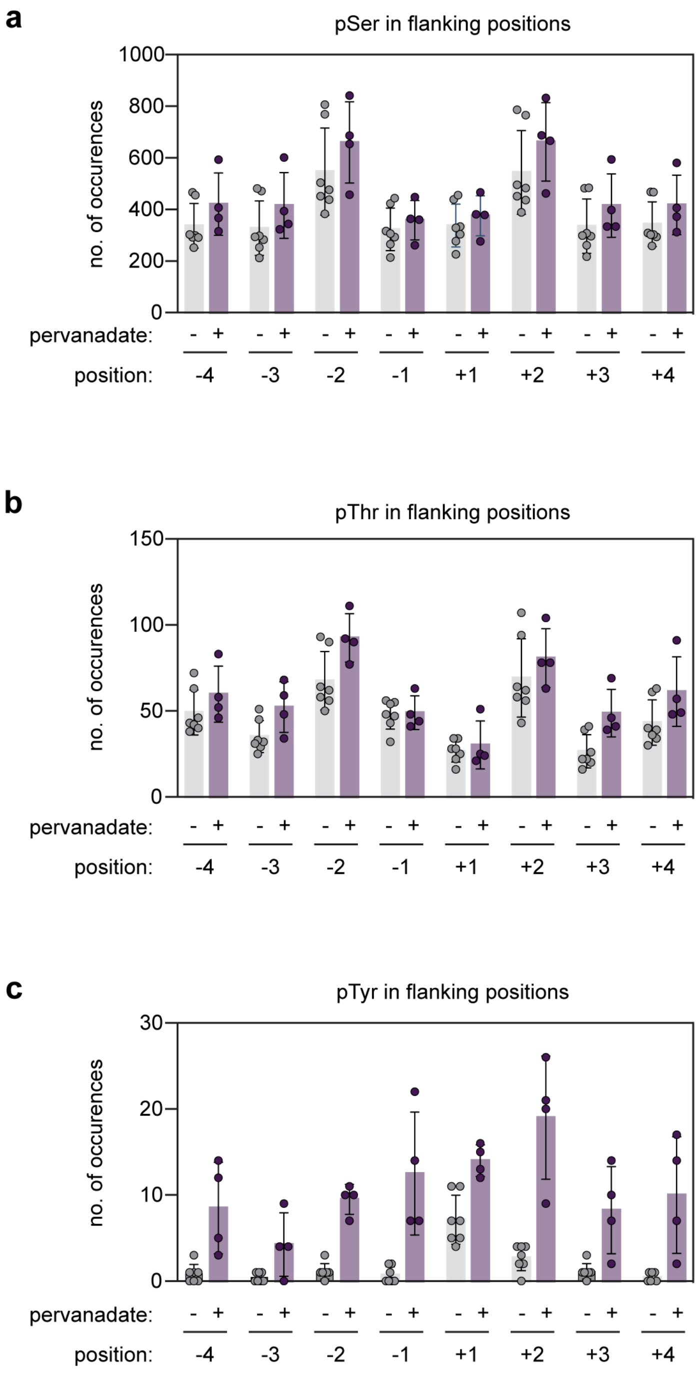

**Figure S6. PTP1B$_{1\text{-}321}$ heatmap timecourses.** a) Timecourse in which all pSer, pThr, and pTyr sites were analyzed. b) Timecourse in which data were filtered to analyze only pSer and pThr sites. No enrichment or depletion in flanking residues is detected because pSer and pThr are not substrates of PTP1B$_{1\text{-}321}$. Z-scores were calculated by comparing positional frequencies to the 0 min timepoint using counts summed across n = 3 biological replicates after filtering for the indicated phosphosite type. Residue-position combinations with Benjamini-Hochberg FDR-adjusted p-values < 0.0001 were considered significant and are outlined in black.

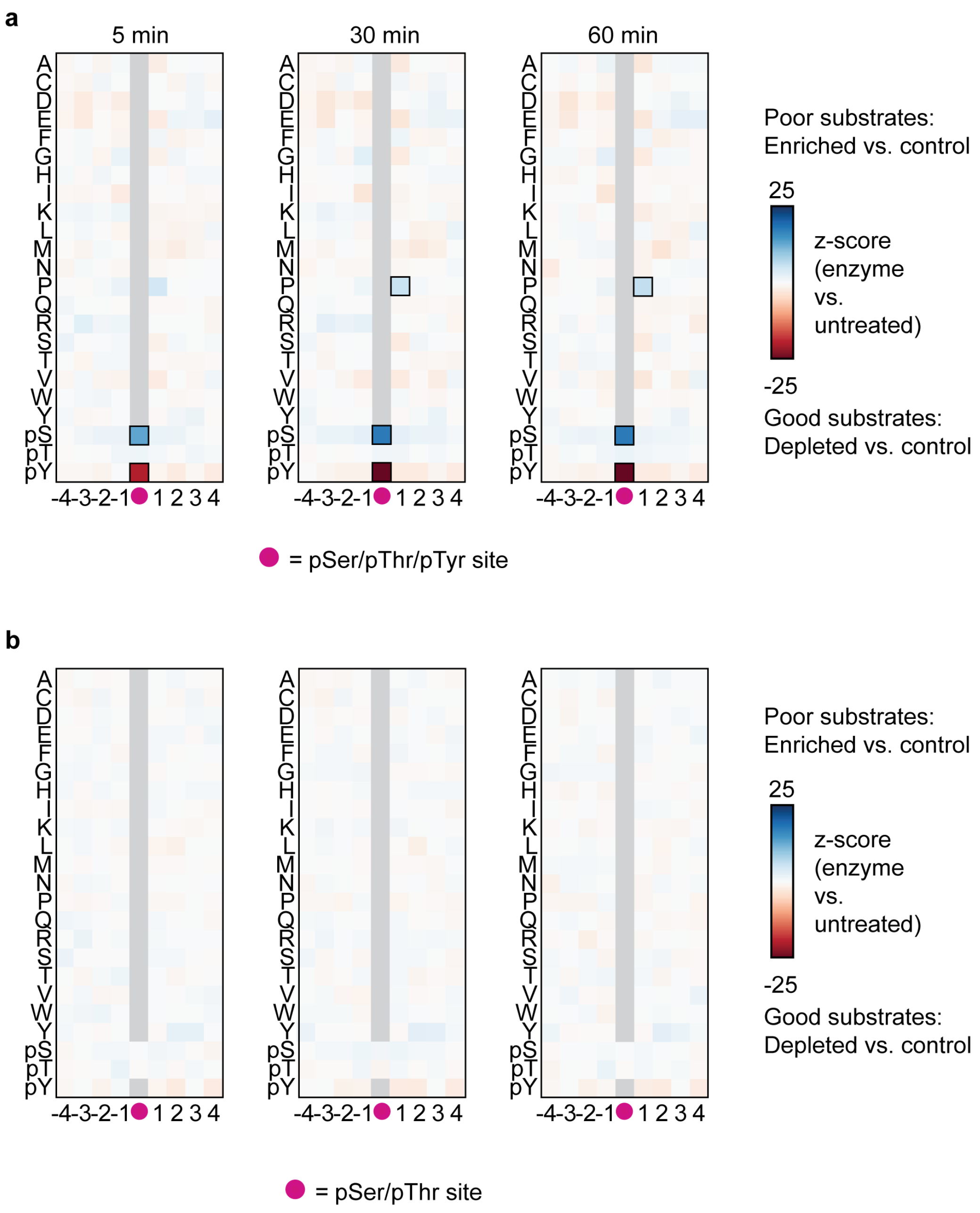

**Figure S7. PP2Ac heatmap timecourses.** a) Timecourse in which data were filtered to analyze only pSer sites. b) Timecourse in which data were filtered to analyze only pThr sites. c) Timecourse in which data were filtered to analyze only pTyr sites. Z-scores were calculated by comparing positional frequencies to the 0 min timepoint using counts summed across n = 3 biological replicates after filtering for the indicated phosphosite type. Residue-position combinations with Benjamini-Hochberg FDR-adjusted p-values < 0.0001 were considered significant and are outlined in black.

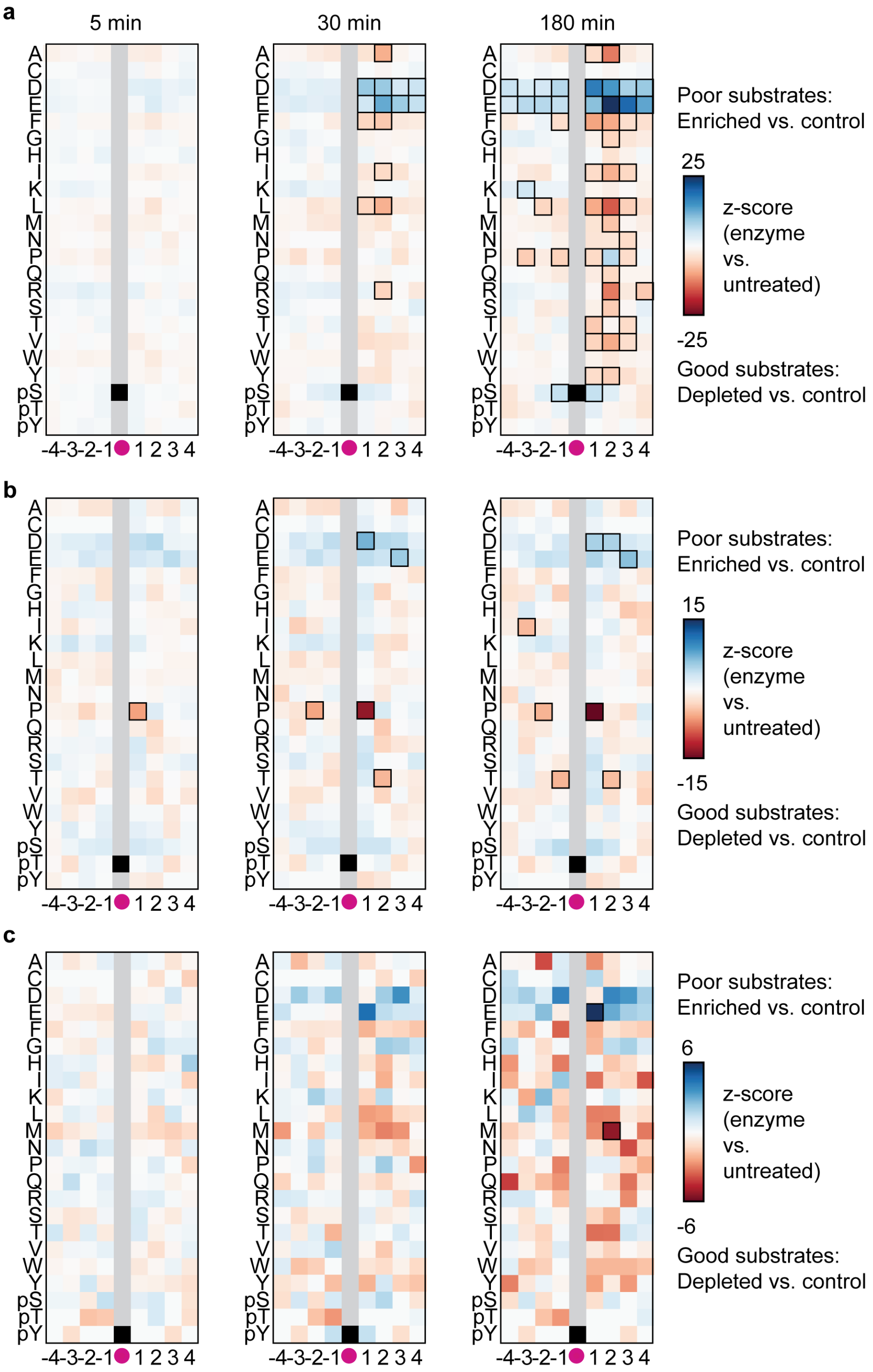

**Figure S8. WipA heatmap timecourses.** a) Timecourse in which all pSer, pThr, and pTyr sites were analyzed. b) Timecourse in which data were filtered to analyze only pSer and pThr sites. Z-scores were calculated by comparing positional frequencies to the 0 min timepoint using counts summed across n = 3 biological replicates after filtering for the indicated phosphosite type. Residue-position combinations with Benjamini-Hochberg FDR-adjusted p-values < 0.0001 were considered significant and are outlined in black.

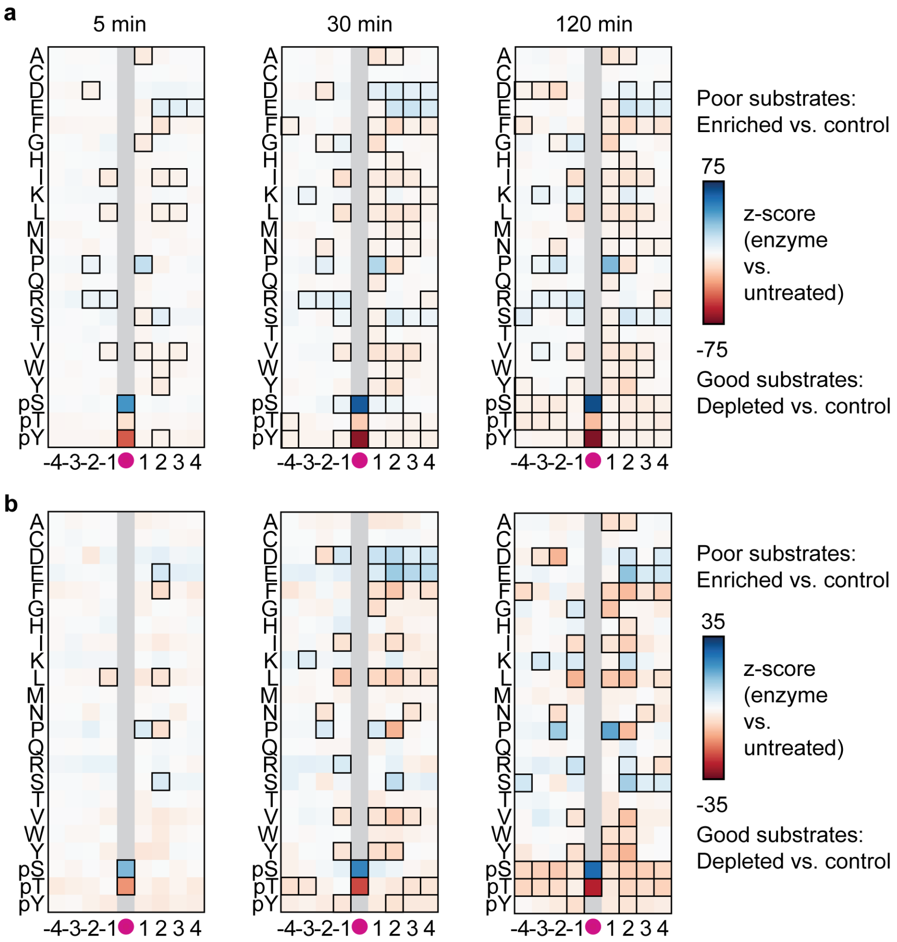

**Figure S9. WipB heatmap timecourses.** a) Timecourse in which all pSer, pThr, and pTyr sites were analyzed. b) Timecourse in which data were filtered to analyze only pSer and pThr sites. c) Timecourse in which data were filtered to analyze only pTyr sites. Z-scores were calculated by comparing positional frequencies to the 0 min timepoint using counts summed across n = 3 biological replicates after filtering for the indicated phosphosite type. Residue-position combinations with Benjamini-Hochberg FDR-adjusted p-values < 0.0001 were considered significant and are outlined in black.

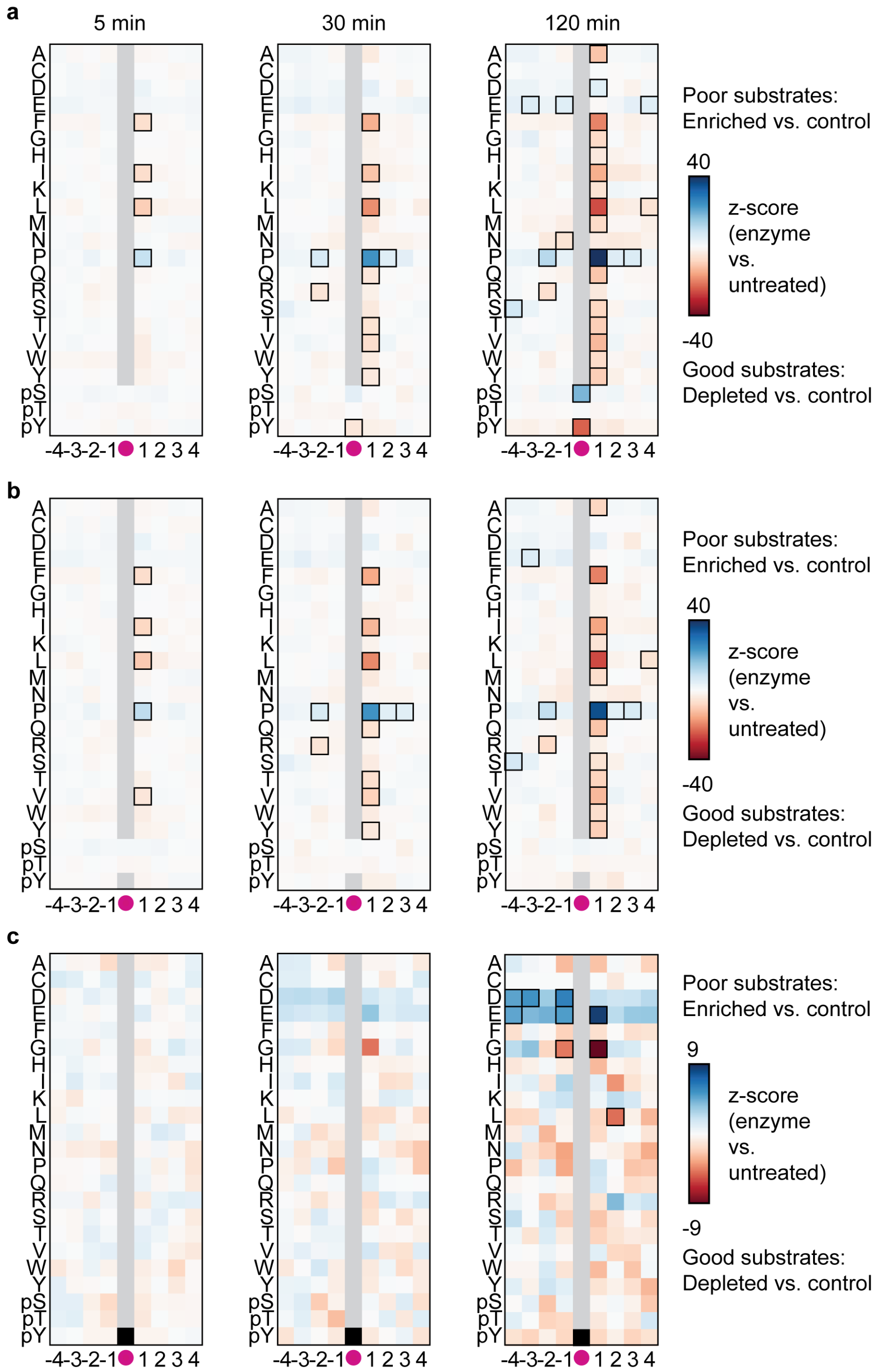

**Figure S10. Distribution of ptmRS site localization scores for β-eliminated sites in OspF-treated PhosPropels from pervanadate-treated HEK293T cells.** Scores were computed using the IMP-ptmRS node in Proteome Discoverer 2.4 (ThermoFisher Scientific). Only sites with scores >90 were included in the analysis.

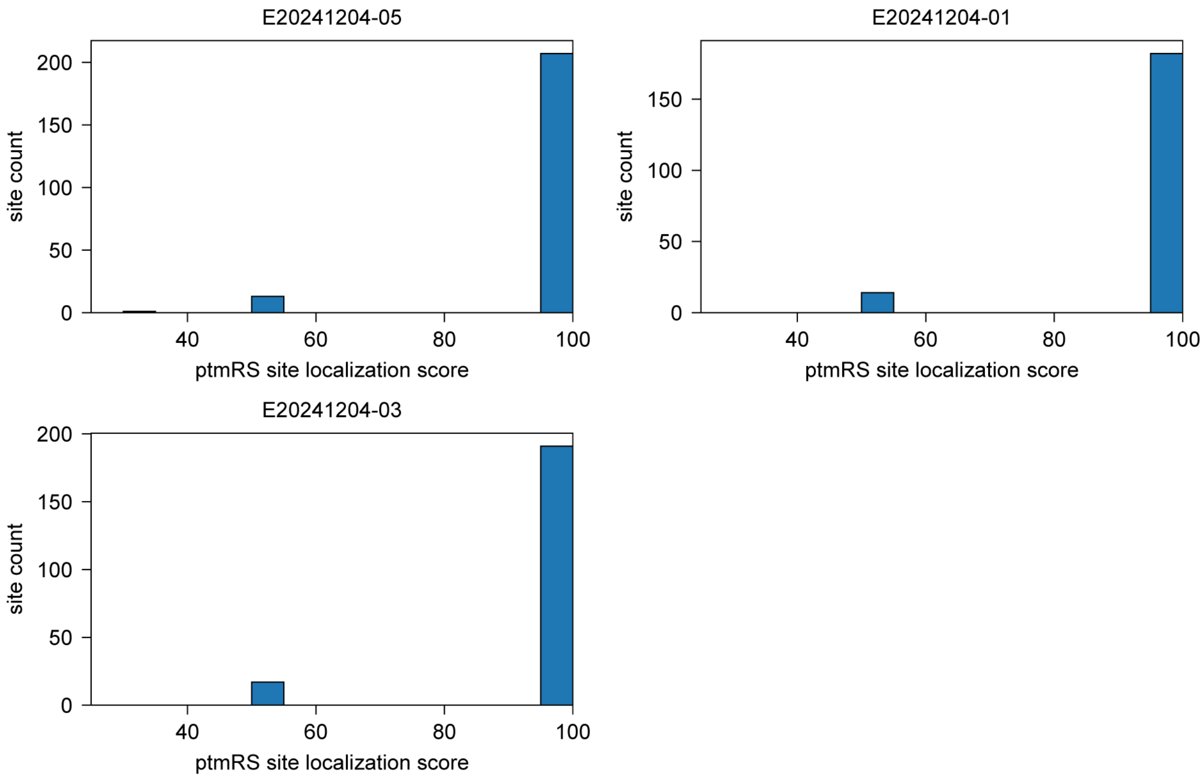

**Figure S11. Characterization of WipA and WipB.** a) SDS-PAGE analysis of purified WipA (lane 2) and WipB (lane 3). b) LC-TOF MS analysis of WipA. The major species has a mass that is -15 Da relative to the expected mass (*). We attribute this to an unexpected modification (e.g., methionine loss + carbamylation) or lower accuracy of the TOF in the higher mass range. c) LC-TOF MS analysis of WipB.

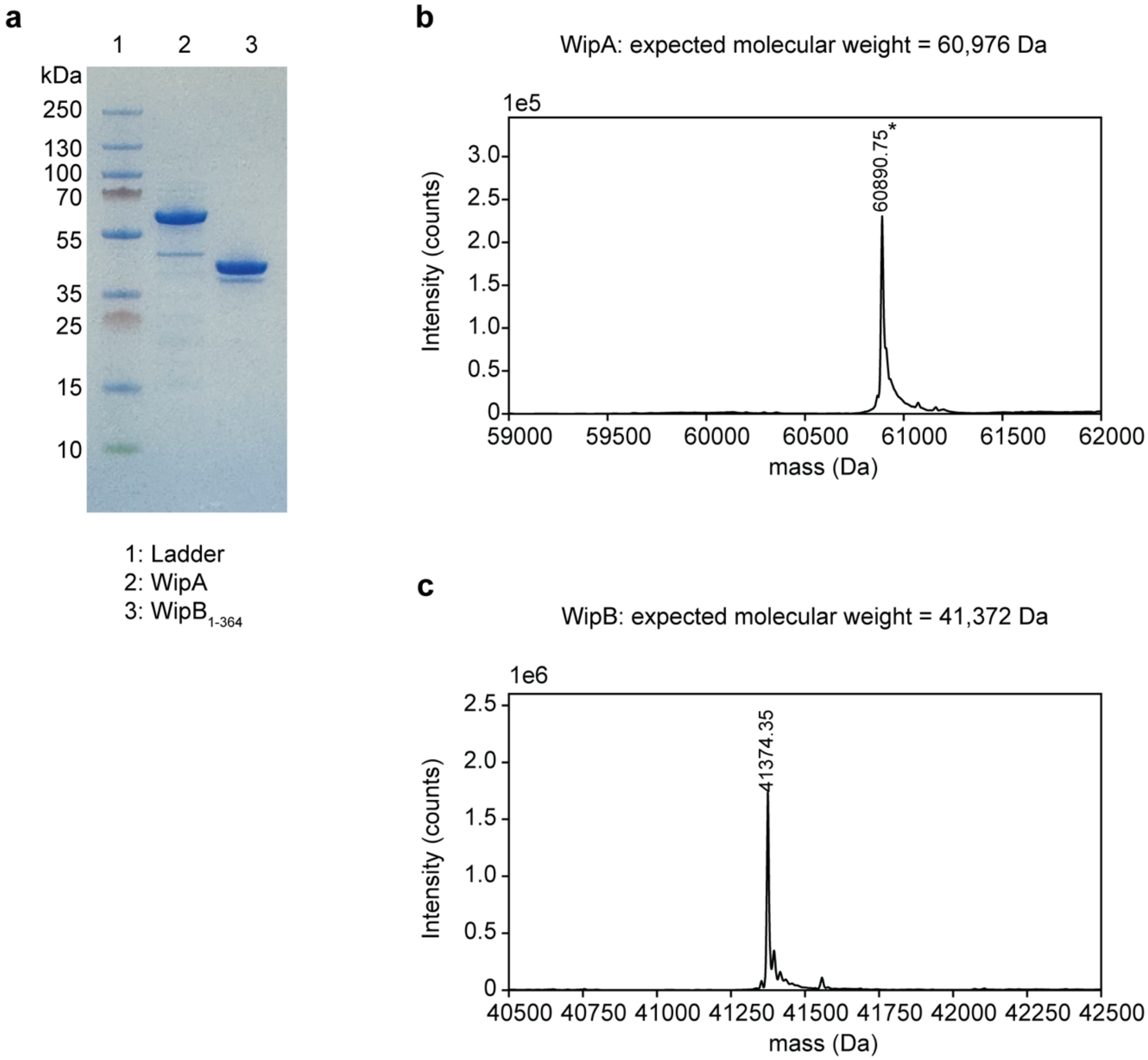

**Figure S12. Characterization of OspF, SpvC, and HopAI.** a) SDS-PAGE analysis of purified OspF (lane 2). b) LC-TOF MS analysis of OspF. c) SDS-PAGE analysis of purified SpvC (lane 2). d) LC-TOF MS analysis of SpvC. e) SDS-PAGE analysis of purified HopAI (lane 2). f) LC-TOF MS analysis of HopAI.

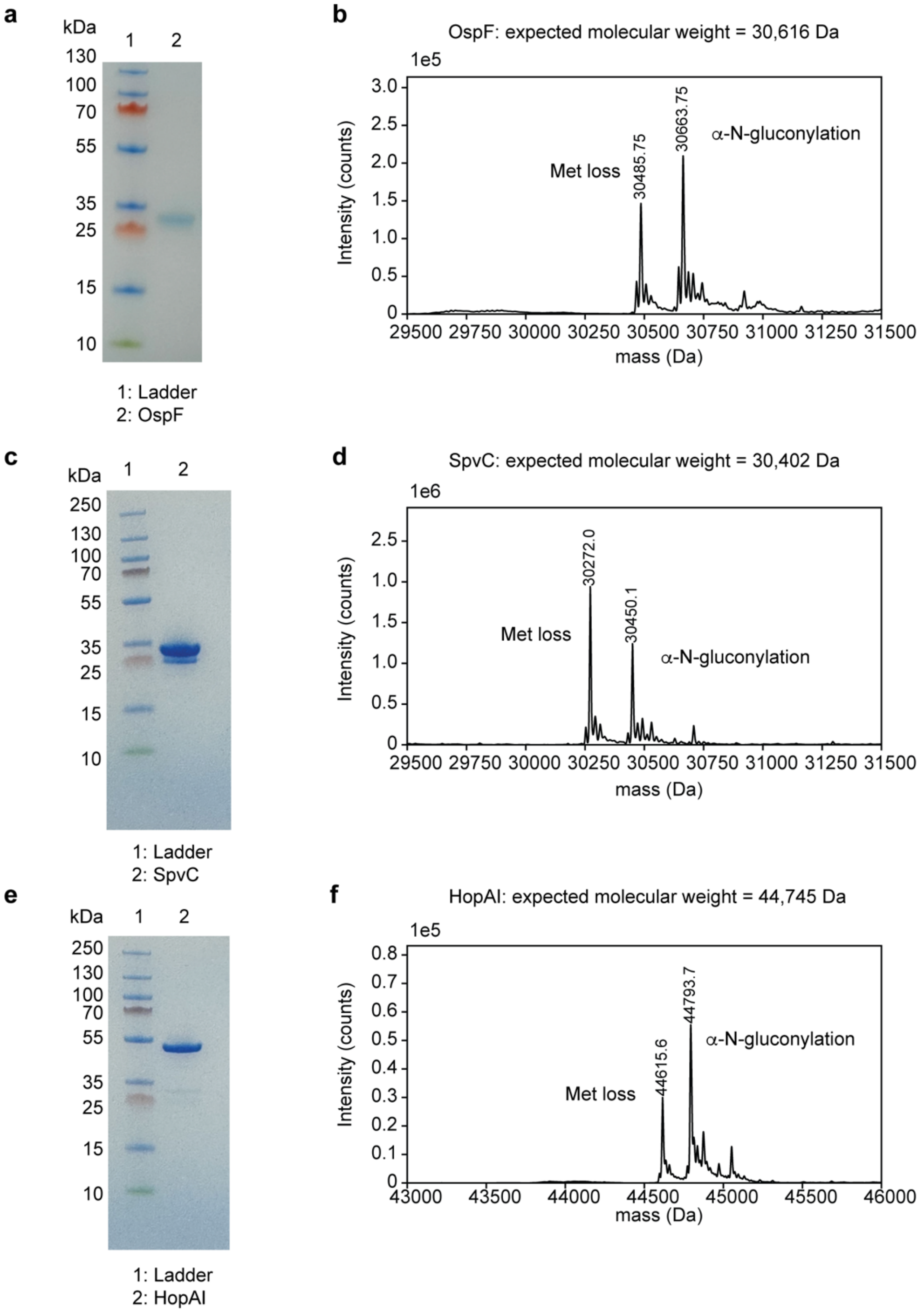

**Supplementary Appendix: Representative spectra for peptides containing dehydroalanine (Dha) and dehydrobutyrine (Dhb) sites.**

**Annotated spectrum for LADFGVAGQL-Dhb-DTQIK.** (A) Spectrum for monoisotopic m/z = 829.94029, z = 2. Matched y-ions are shown in blue and matched b-ions are shown in red. Ammonia loss peaks are marked with a * and water loss peaks are marked with a •. Ions $y_4$, $y_6$, $y_7$, $y_8$, $y_9$, $y_{10}$, $y_{11}$, $y_{12}$, and $y_{13}$ (bolded) are shifted by -18.0 Da relative to the theoretical m/z for the unmodified peptide, supporting localization of the modification at peptide position 13 (ptmRS site probability = 100%). (B) Theoretical m/z for y-ion and b-ion series of unmodified LADFGVAGQLTDTQIK and LADFGVAGQLTDTQIK in which Thr 13 has been converted to Dhb. Matched y-ions are colored in blue and matched b-ions are colored in red. Fragments with shifted m/z between the modified and unmodified peptides are bolded.

A

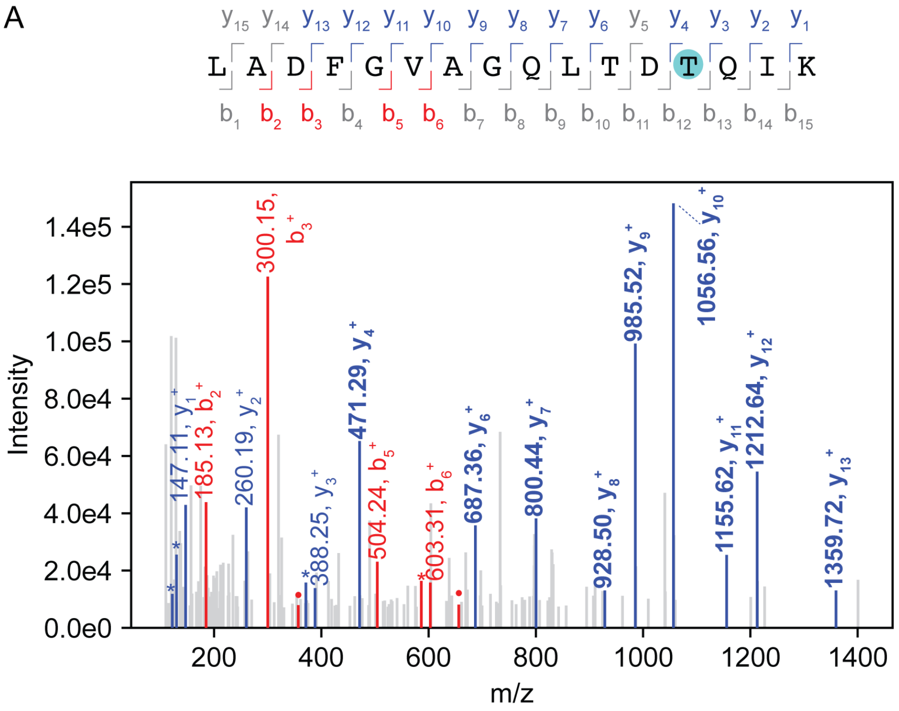


B

| b⁺, unmodified | b⁺, modified | b-ion # | | y-ion # | y⁺, unmodified | y⁺, modified |
|---|---|---|---|---|---|---|
| 114.09134 | 114.09134 | 1 | L | 16 | | |
| 185.128454 | 185.128454 | 2 | A | 15 | 1563.801313 | 1545.790748 |
| 300.155397 | 300.155397 | 3 | D | 14 | 1492.7642 | 1474.753635 |
| 447.223811 | 447.223811 | 4 | F | 13 | 1377.737257 | **1359.726692** |
| 504.245275 | 504.245275 | 5 | G | 12 | 1230.668843 | **1212.658278** |
| 603.313689 | 603.313689 | 6 | V | 11 | 1173.647379 | **1155.636814** |
| 674.350803 | 674.350803 | 7 | A | 10 | 1074.578965 | **1056.5684** |
| 731.372267 | 731.372267 | 8 | G | 9 | 1003.541851 | **985.531286** |
| 859.430844 | 859.430844 | 9 | Q | 8 | 946.520387 | **928.509822** |
| 972.514908 | 972.514908 | 10 | L | 7 | 818.46181 | **800.451245** |
| 1073.562587 | 1073.562587 | 11 | T | 6 | 705.377746 | **687.367181** |
| 1188.58953 | 1188.58953 | 12 | D | 5 | 604.330067 | 586.319502 |
| 1289.637208 | 1271.626643 | 13 | T | 4 | 489.303124 | **471.292559** |
| 1417.695786 | 1399.685221 | 14 | Q | 3 | 388.255446 | 388.255446 |
| 1530.77985 | 1512.769285 | 15 | I | 2 | 260.196868 | 260.196868 |
| | | 16 | K | 1 | 147.112804 | 147.112804 |

**Annotated spectrum for TPEELDD-Dha-DFETEDFDVR.** (A) Spectrum for monoisotopic m/z = 1070.95561, z = 2. Matched y-ions are shown in blue and matched b-ions are shown in red. Ammonia loss peaks are marked with a * and water loss peaks are marked with a •. Ions $y_{11}$-$y_{16}$, and $b_8$, $b_9$, and $b_{10}$ (bolded) are shifted by -18.0 Da relative to the theoretical m/z for the unmodified peptide, supporting localization of the modification at peptide position 8 (ptmRS site probability = 100%). (B) Theoretical m/z for y-ion and b-ion series of unmodified TPEELDDSDFETEDFDVR and TPEELDDSDFETEDFDVR in which Ser 8 has been converted to Dha. Matched y-ions are colored in blue and matched b-ions are colored in red. Fragments with shifted m/z between the modified and unmodified peptides are bolded.

A

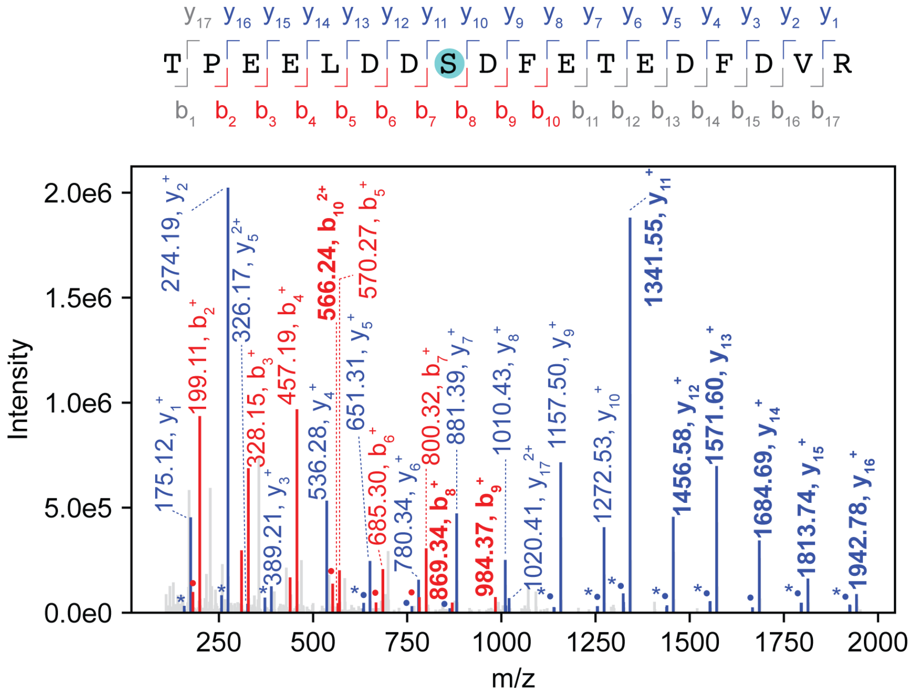


B

| b⁺, unmodified | b⁺, modified | b-ion # | | y-ion # | y⁺, unmodified | y⁺, modified |
|---|---|---|---|---|---|---|
| 102.055 | 102.055 | 1 | T | 18 | | |
| 199.1077 | 199.1077 | 2 | P | 17 | 2057.846 | 2039.835 |
| 328.1503 | 328.1503 | 3 | E | 16 | 1960.793 | **1942.782** |
| 457.1929 | 457.1929 | 4 | E | 15 | 1831.75 | **1813.74** |
| 570.277 | 570.277 | 5 | L | 14 | 1702.708 | **1684.697** |
| 685.3039 | 685.3039 | 6 | D | 13 | 1589.624 | **1571.613** |
| 800.3309 | 800.3309 | 7 | D | 12 | 1474.597 | **1456.586** |
| **887.3629** | 869.3523 | 8 | S | 11 | 1359.57 | **1341.559** |
| **1002.39** | 984.3793 | 9 | D | 10 | 1272.538 | 1272.538 |
| **1149.458** | 1131.448 | 10 | F | 9 | 1157.511 | 1157.511 |
| 1278.501 | 1260.49 | 11 | E | 8 | 1010.443 | 1010.443 |
| 1379.549 | 1361.538 | 12 | T | 7 | 881.3999 | 881.3999 |
| 1508.591 | 1490.581 | 13 | E | 6 | 780.3523 | 780.3523 |
| 1623.618 | 1605.607 | 14 | D | 5 | 651.3097 | 651.3097 |
| 1770.686 | 1752.676 | 15 | F | 4 | 536.2827 | 536.2827 |
| 1885.713 | 1867.703 | 16 | D | 3 | 389.2143 | 389.2143 |
| 1984.782 | 1966.771 | 17 | V | 2 | 274.1874 | 274.1874 |
| | | 18 | R | 1 | 175.119 | 175.119 |

**Annotated spectrum for HTDDEM-Dhb-G-pTyr-VATR.** (A) Spectrum for monoisotopic m/z = 779.30698, z = 2. Matched y-ions are shown in blue and matched b-ions are shown in red. Ammonia loss peaks are marked with a * and water loss peaks are marked with a •. Ions $y_5$ and $y_6$ are shifted by 80 Da relative to the theoretical m/z for the unmodified peptide, supporting localization of the phospho modification at peptide position 9 (ptmRS site probability = 99.5%). Ions $b_7$ and $b_8$ are shifted by -18 Da relative to the theoretical m/z for the unmodified peptide, supporting localization of the dehydration modification at position 7. Ions $y_7$-$y_{12}$ and $b_9$ are shifted by 62 Da (80 Da for phospho at position 9 and -18 Da for dehydration on position 7) relative to the theoretical m/z values for the unmodified peptide, supporting localization the dehydration modification at peptide position 7 (ptmRS site probability = 100%). (B) Theoretical m/z for y-ion and b-ion series of HTDDEM-Dhb-G-pTyr-VATR. Matched y-ions are colored in blue and matched b-ions are colored in red. Ions in bold have the dehydration modification, ions in italic have the phosphor modification, and ions in bold italic have both modifications.

A

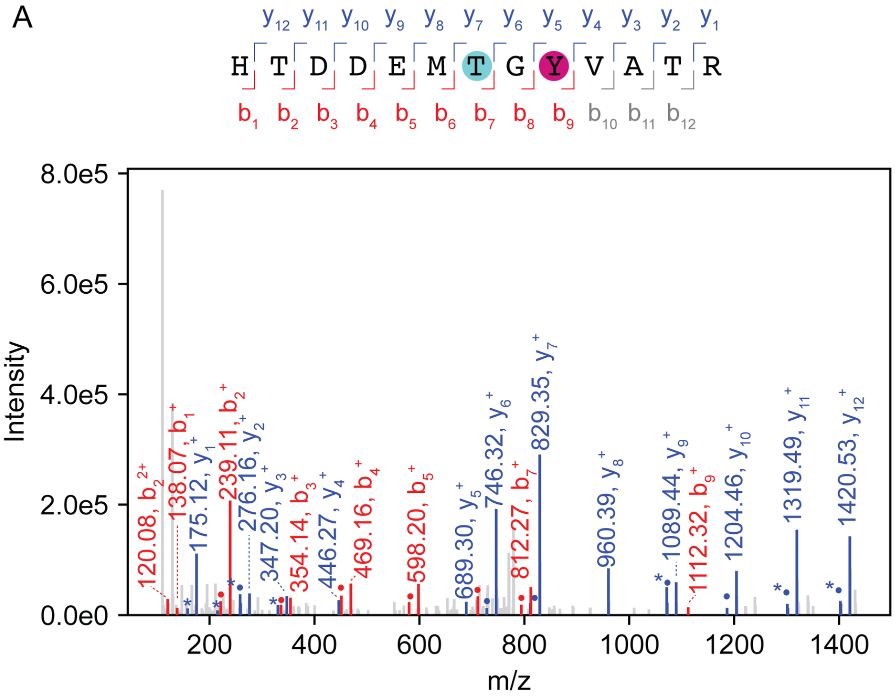


B

| b⁺ | b-ion # | | y-ion # | y⁺ |
|---|---|---|---|---|
| 138.0662 | 1 | H | 13 | |
| 239.1139 | 2 | T | 12 | ***1420.545*** |
| 354.1408 | 3 | D | 11 | ***1319.497*** |
| 469.1678 | 4 | D | 10 | ***1204.47*** |
| 598.2103 | 5 | E | 9 | ***1089.443*** |
| 729.2508 | 6 | M | 8 | ***960.4009*** |
| **812.2879** | 7 | T | 7 | ***829.3604*** |
| **869.3094** | 8 | G | 6 | *746.3233* |
| ***1112.339*** | 9 | Y | 5 | *689.3018* |
| 1211.407 | 10 | V | 4 | 446.2722 |
| 1282.445 | 11 | A | 3 | 347.2037 |
| 1383.492 | 12 | T | 2 | 276.1666 |
| | 13 | R | 1 | 175.119 |

**Supplementary Note: Statistical framework for phosphoeraser specificity profiling using PhosPropels.**

The PhosPropel approach enables statistical comparison of amino acid frequencies at positions surrounding a central modified site in an enzyme-treated sample versus an appropriate control. Here, we describe general considerations that make this comparison possible and the specific analyses that were performed in our manuscript.

**Background set**

Rather than relying on a theoretical or computationally derived background set, we used an experimentally measured control as the background for all statistical comparisons. This approach ensures that enrichment or depletion is assessed relative to the actual position-specific distribution of amino acids present in the phosphoproteome-derived peptide library. The experimental background set accounts for position-specific biases in amino acid composition surrounding phosphosites (e.g., Pro occurs much more frequently in the position following pSer than its overall frequency in the proteome) and for experimental treatments that alter phosphosite composition in the library (e.g. treatment with pervanadate or other phosphatase inhibitors).

**Sample set**

Sample sets are generated by treating the phosphoproteome-derived peptide library with a phosphoeraser enzyme of interest. To profile phosphatase specificity, the remaining phosphosites in the library can be analyzed and the position-specific frequencies of each amino acid in each position surrounding the phosphosites can be calculated. Good substrates contain features that are depleted from the library, while poor substrates contain features that persist in library after enzyme treatment. To profile phospholyase specificity, the appearance of β-eliminated Ser and Thr residues (dSer and dThr) can be analyzed and the position-specific frequency of each amino acid in each position surrounding the β-eliminated site can be calculated. Good substrates contain features that are enriched in positions surrounding the β-eliminated sites, while poor substrates contain features that appear less frequently surrounding β-eliminated sites.

**Statistical comparison to identify sequence features that influence phosphoeraser substrate preference**

In comparing the background set and the sample set, the null hypothesis is that the frequency of amino acid X at position i is the sample in the treated sample versus background sample, and that all amino acids at position i are equally likely to be found among substrates of the enzyme under study. The alternative hypothesis is that the frequency of amino acid X at position i is different in the treated sample versus background sample, reflecting enzyme-specific substrate preferences.

To test these hypotheses, we calculated z-scores for each amino acid-position pair according to the following formula:

$$z = \frac{p_{sample} - p_{control}}{SE}$$

where $p_{sample}$ is the observed frequency of amino acid X at position i in the enzyme-treated sample, $p_{control}$ is the observed frequency of amino acid X at position i in the background sample, and SE is calculated according to the formula:

$$SE = \sqrt{\frac{p_{sample}(1 - p_{sample})}{n_{sample}} + \frac{p_{control}(1 - p_{control})}{n_{control}}}.$$

We treated the frequency of each amino acid at each position as a binomial proportion, assuming that each peptide provides an independent observation of residue identity at a given position. Under this assumption, the variance of each frequency estimate can be calculated based on the binominal distribution, and differences in frequency can be standardized using the combined standard error. We applied the normal approximation to the binomial distribution for z-score calculation based on the central limit theorem. This approximation is appropriate given the large number of measurements in both conditions and the sufficient count of most amino acid-position combinations.

The z-score reflects how many standard errors the observed difference is from the null expectation (i.e., the background frequency). The z-score approach accounts for not only the difference in observed frequencies, but also for variability associated with the number of observations of each amino acid in each position due to uneven amino acid usage in the proteome and due to experimental considerations. Because z-scores incorporate the number of observations for each amino acid at each position, they effectively scale the confidence of each comparison. For example, a large difference in frequency between the treated sample and the untreated control that is supported by a large number of observations produces a higher-magnitude z-score, while differences based on only a few measurements produce z-scores of lower magnitudes.

We note that our statistical approach is very similar to that used to produce IceLogos with small changes in how SE is calculated based on our experimental design.

**Analysis of position-specific frequencies in PhosPropels compared to the proteome**

For analysis of the phosphosite-flanking residues in PhosPropels, we chose as our sample set all phosphosites confidently identified and localized using LC/MS analysis. We aligned these phosphosites and counted the frequency of each amino acid at each position four residues N-terminal (- side) and four residues C-terminal (+ side) to the phosphosite. As a control/background sample, we measured the overall frequency of each of the amino acid in the sample in a manner that was not position-specific. We favored this approach as opposed to using previously reported information about the abundance of each amino acid in the human as it allowed us to measure the frequency of occurrence of pSer, pThr, and pTyr in the sample, enabling us to ask whether these amino acids were enriched or depleted relative to other phosphosites.

**PhosPropel-based profiling of phosphatase specificity**

For analysis of phosphatase specificity using PhosPropels, we chose as our sample set all phosphosites confidently identified and localized using LC-MS/MS analysis. We chose as our control/background set all phosphosites confidently identified and localized using LC-MS/MS in an untreated or 0 min timepoint sample. To compare the distributions of phosphosite features, we aligned the phosphosites in both sample and control, computed the position-specific frequencies of each amino acid, and calculated z-scores as described above to compare them. We reasoned that features associated with good phosphatase substrates would be specifically depleted from the library upon phosphatase treatment, while features associated with poor substrates would become enriched in the library upon phosphatase treatment.

**PhosPropel-based profiling of phospholyase specificity**

For analysis of phospholyase specificity using PhosPropels, we chose as our sample set all β-eliminated sites that were confidently identified and localized using LC-MS/MS analysis. These sites could be identified based on a -18.01 Da mass shift relative to the unmodified amino acid. We chose as our control/background set all phosphosites confidently identified and localized using LC-MS/MS in the same sample. We aligned the β-eliminated sites in the sample and the phosphosites in the control, computed the position-specific frequencies of each amino acid, and calculated z-scores as described above to compare them. We reasoned that features associated with good phospholyase substrates would be specifically enriched among β-eliminated sites, while features associated with poor substrates would become enriched among phosphosites.

**Comparison of wild-type OspF vs. variant OspF specificity**

For comparison of the sequence specificity of wild-type OspF and OspF variants, we chose as our sample set all β-eliminated sites that were confidently identified and localized using LC-MS/MS analysis in the variant-treated sample. We chose as our control/background set all β-eliminated sites confidently identified and localized using LC-MS/MS in a wild-type OspF-treated sample. We aligned the β-eliminated sites in the sample and in the control, computed the position-specific frequencies of each amino acid, and calculated z-scores as described above to compare them. Features associated with improved variant activity compared to wild-type were specifically enriched in among β-eliminated sites in the variant-treated sample, while features associated with poor variant substrates substrates were specifically enriched among β-eliminated sites in the wild-type OspF-treated control.
