## Supplementary Dataset 6 - HEK293T pervanadate-treated phosphopeptide library treated with lambda phosphatase for "Phosphoproteome-derived peptide libraries for deep specificity profiling of phosphatases and phospholyases": README.txt

Title: Phosphoproteome-derived peptide libraries for phosphatase specificity profiling

Authors: Katarzyna Radziwon, Laura Campbell, Lauren E. Mazurkiewicz, Sopo Jalalishvili, Izabelle Eppinger, Aanika Parikh, and Amy M. Weeks
Institution: University of Wisconsin--Madison
Contact:

Description:
This dataset accompanies the manuscript "Phosphoproteome-derived peptide libraries for deep specificity profiling of enzymatic phosphorylation erasers". This dataset contains csv files containing peptide identification search results for replicate phosphoproteome-derived peptide libraries made from HEK293T cells that were treated with lambda phosphatase and analyzed at different time points.

Data collection and analysis:
- Phosphoproteome-derived peptide libaries were prepared using TiO2 enrichment as described in the accompanying manuscript.
-Libraries were treated with lambda phosphatase as described in the manuscript and quenched at different time points
- LC-MS/MS data were collected on an Orbitrap Exploris 480 (ThermoFisher Scientific)
- Search: Database search was performed with Proteome Discoverer v2.4 using SEQUEST HT with Percolator post-processing
- Site localization: Phosphosite localization was scored using ptmRS
- Database: SwissProt human (downloaded January 24, 2020)


How the data are organized:

README.txt — This file

Subdirectories - replicate datasets for each timepoint are arranged into the following subdirectories:
├── 0min/
├── 5min/
├── 30min/
└── 180min/

Each folder contains csv files corresponding to the Peptide Groups tab of search results from Proteome Discoverer. Each csv file is one replicate for the timepoint named in the subdirectory name. The columns in each csv are:

Checked
Confidence
Annotated Sequence - this column is used in downstream analysis to extract the peptide sequence
Modifications - this column contains information about the localization of phosphorylation, dehydration, and other modifications. An example is '1xPhospho [S17(98.6)]', which describes a peptide with one phosphosite at positions S17 with a ptmRS site localization score of 98.6
Qvality PEP
Qvality q-value
# Protein Groups
# Proteins
# PSMs
Master Protein Accessions - Uniprot accession numbers
Positions in Master Proteins
Modifications in Master Proteins - this column contains information about the location of modifications within the protein from which the peptide is derived. An example is 'P05387 1xPhospho [S17(98.6)]', where P05387 is the protein that this peptide is derived from, the peptide has one phosphorylation site at position S17 in the protein sequence, and the ptmRS site localization score is 98.6
# Missed Cleavages
Theo. MH+ [Da]
Found in Sample: [S10] F10: Sample
Confidence (by Search Engine): Sequest HT
Percolator q-Value (by Search Engine): Sequest HT
Percolator PEP (by Search Engine): Sequest HT
XCorr (by Search Engine): Sequest HT
